# Plumage manipulation alters the integration of social behavior, physiology, internal microbiome, and fitness

**DOI:** 10.1101/826719

**Authors:** Conor C. Taff, Cedric Zimmer, David Scheck, Thomas A. Ryan, Jennifer L. Houtz, Melanie R. Smee, Tory A. Hendry, Maren N. Vitousek

## Abstract

Signals often covary with physiological and behavioral traits to form an axis of integrated phenotypic variation associated with reproductive performance. This pattern of phenotypic integration could result from intrinsic between-individual differences that are causally related to signal production, physiology, and behavior. Alternatively, signal expression itself might generate dynamic feedback between physiology, behavior, and the experienced social environment, resulting in an integrated phenotype. Here, we manipulated the plumage of female tree swallows (*Tachycineta bicolor*) to decouple the expression of a social signal from any pre-existing behavioral or physiological differences. We collected a time series of physiological samples, monitored social interactions with a sensor network, and tracked reproductive performance. Relative to sham controls, dulled females experienced an altered social environment; overall, these females were visited more by conspecific females and less by males. Dulled females subsequently changed their own behavior by initiating fewer interactions and increasing nestling provisioning. These differences resulted in an altered internal microbiome and glucose levels and, ultimately, dulled females produced more offspring. Moreover, dulled females produced larger clutches than control females in the year after the manipulation. Thus, signal variation alone—independent from any pre-existing differences—had a sustained causal affect on a suite of integrated traits. This finding suggests that dynamic feedback may play an important role in coordinating an integrated signaling phenotype. Our results have implications for understanding how variation in signal expression arises and is maintained and the extent to which the information encoded in signals is contingent upon their use in a social environment.

## INTRODUCTION

Animal signals are often one part of an integrated suite of phenotypic traits that includes a variety of physiological and behavioral measures (e.g. Levin et al. 2016). Individuals that are best able to coordinate signals, behavior, and physiology are likely to achieve the highest relative fitness. As a result, mate choice based on signals often results in selection of mates that also have favorable behavioral and physiological phenotypes. One long-standing line of research into the evolution of signals has focused on identifying mechanism(s) that could generate and maintain this apparent honesty in signal variation. A large number of potential mechanisms based on trade-offs or limited resources have been proposed over the last 30 years. For example, signal honesty might be enforced by early nutritional environment (Nowicki et al. 2002), the ability to tolerate high hormone titers (Folstad & Karter 1992), a trade-off between investing in signals and oxidative defense (Alonso-Alvarez et al. 2008), or mitonuclear compatibility (Hill et al. 2019), among other possibilities.

While these proposed mechanisms differ in their details, a key similarity is that they are all commonly described and tested as unidirectional causal paths. That is, intrinsic variation associated with condition, experience, or genotype directly contributes to signal development, which in turn influences performance in the social environment, resulting in fitness differences. However, it has also long been known that possessing a certain signaling phenotype can change the social environment that animals experience (Chaine et al. 2018; Webster et al. 2017) and that social experience can alter subsequent physiology (Liu et al. 1997), behavior (Cornelius et al. 2018; Hirschenhauser & Oliveira 2006), and signal development (Dey et al. 2014; Maia et al. 2012). As a consequence, manipulating signals can result in altered physiology, including alterations to measures that may be related to signal production in the first place (Levin et al. 2018; Safran et al. 2008; Tibbetts et al. 2016; Vitousek et al. 2013). While both classic honesty mechanisms and dynamic feedback might produce similarly coordinated signaling phenotypes, the underlying processes are quite different (Rubenstein & Hauber 2008; Tibbetts 2014; Vitousek et al. 2014). Critically, in the dynamic feedback scenario, phenotypic integration emerges as the result of regulatory processes linking physiology, behavior, the social environment, and signal production rather than as the end product of a unidirectional causal chain. Distinguishing between these two possibilities remains challenging both because they may result in similar patterns of trait correlation and because convincingly demonstrating dynamic feedback requires a time series of physiological data coupled with detailed behavioral observations and measures of performance.

Here, we studied the causal role that signals play in integrating physiology, behavior, internal microbiome composition, and reproductive success in tree swallows (*Tachycineta bicolor*). Previous work in this population demonstrated that brighter white breast plumage in female tree swallows is associated with increased resilience to challenging conditions, greater social interactivity, a stronger glucocorticoid stress response, and genome wide DNA methylation patterns (Taff et al. 2019a; Taff et al. 2019b). Thus, plumage brightness is part of an integrated suite of traits typically associated with increased performance. In this study, we asked to what extent these correlations arise as a direct result of conspecific behavioral responses to signal expression (dynamic integration hypothesis). Alternatively, the correlations that we previously observed might be the consequence of intrinsic between individual differences that directly influence signals, physiology, and behavior independently (fixed integration hypothesis). In reality, these two hypotheses are not mutually exclusive and the suite of traits correlated with plumage may result from a combination of dynamic and fixed causes.

We experimentally dulled the white plumage of female tree swallows to decouple signal expression from any pre-existing differences in behavior and physiology. We then used a remote monitoring network to record detailed behavioral data for the entire breeding population. We combined this behavioral data with repeated measurements of corticosterone, glucose, and internal microbiome diversity. Using these measures, we compared the behavior, physiology, and reproductive success of dulled females to that of sham controls. We assumed that females could not directly assess their own signaling phenotype, thus any downstream effects of plumage manipulation on focal female physiology and behavior must represent a causal impact of signal expression. We did not make specific predictions about how signal manipulation would impact individual traits. Rather, we predicted that if the suite of traits associated with plumage were integrated due—in part—to the social response of conspecifics to signal expression then we would detect downstream impacts of plumage dulling on conspecific behavior, resulting in subsequent changes in focal female behavior, physiology, and reproductive success. A failure to detect treatment differences would suggest that the correlations that we observed previously are explained by intrinsic between individual differences rather than causal effects of signal expression during the breeding season *per se*. After testing for an overall effect of dulling, we explore several plausible mechanisms that could link signal expression to performance.

## METHODS

### General Field Methods

We studied breeding tree swallows at Cornell’s Experimental Ponds in Ithaca, New York from April to July 2017 (42.503° N, 76.437° W). Tree swallows at these sites have been monitored continuously since 1986 and we followed a standardized protocol for recording breeding activity at the site (for extended details on field methodology see Vitousek et al. 2018a; Winkler et al. 2013).

The nest boxes used for this study are arranged in two large grids around artificial ponds; the two grids are separated by 2 km of woods and boxes within each grid are spaced 20 meters apart. We checked nest boxes every other day beginning in late April and recorded key breeding dates for each pair (clutch initiation date, clutch completion date, and fledging or failure date). Around the expected hatching date, we checked nests every day so that exact hatch dates could be determined. For this study, we included 70 nests (36 control and 34 manipulated females; see below). These nests produced a total of 372 eggs and 228 nestlings that survived to be sampled on day 12 (see below).

At each nest, females were captured three times during the breeding season; these three captures were scheduled on day 6-7 of incubation, day 3-4 after hatching, and day 7-8 after hatching. All adult female captures occurred between 7 am and 10 am to minimize the effect of circadian variation in the physiological parameters that we planned to measure (see below). Males were captured once on day 3-8 after hatching. Because males are more difficult to capture and because our experiment was focused on females, we only attempted to capture males on a maximum of two days to minimize disturbance at the nest box. We also were unable to catch males at any nest that failed before day 3 after hatching (*n* = 6). In total, males were sampled at 46 out of 70 nests included in this study. For nestlings, we took a total brood mass measurement on day 6 (all nestlings were weighed together but not individually marked) and then revisited nests on day 12 after hatching for banding and morphological measurements. With this sampling scheme, we had reliable data on total clutch size, number of eggs that hatched, number of chicks alive on day 6 and 12, and number of chicks that successfully fledged.

All adults and nestlings were banded with a unique USGS aluminum band. Adults were also equipped with a passive integrated transponder tag (PIT tag) on the other leg that encoded a unique ten-digit hexadecimal string and could be read by the radio frequency identification (RFID) readers installed on each nest box at the site (see details below). The PIT tags were encased in silicone shrink tubing and glued to a celluloid color band that was sealed with acetone (Vitousek et al. 2018b). We also collected a set of morphological measurements at each capture (head plus bill length, flattened wing cord length, and mass), except for the third female capture, when we only measured mass. At the first and third capture, we collected a cloacal swab to provide a measure of the internal microbiome. For these samples, we cleaned the area around the cloaca with an alcohol wipe to prevent contamination from external bacteria and then inserted a sterile flocked swab 1.5 cm into the cloaca (Puritan Medical Products Company LLC). The swab was removed while slowly rotating (as in Vo & Jedlicka 2014) and stored in 1 mL of RNAlater in a sterile microcentrifuge tube at −80° C until DNA extraction (see below).

For adults, we collected 6-8 feathers from the center of the white breast to measure breast brightness (initial sample) and to assess the effects of treatments (subsequent samples). For females, we recorded age as second year (SY) if females possessed the brown dorsal plumage that is characteristic of SY tree swallows or after-second year (ASY) if the dorsal plumage was partially or all blue-green. Males cannot be reliably aged based on plumage and were all considered after hatch year.

In addition to the measurements described above, we collected blood samples from females to assess glucose levels, and circulating corticosterone. At the first and second female capture we collected a series of three blood samples by brachial venipuncture. First, a baseline sample (< 70 μl) was collected within 3 minutes of capture. Next, a stress-induced sample (< 30 μl) was collected 30 minutes after capture to measure maximal corticosterone elevation. Finally, we measured variation in the efficacy of negative feedback by injecting birds with dexamethasone (4.5 μl g^−1^; Mylan® 4mg ml^−1^ dexamethasone sodium phosphate, product no.: NDC 67457-422-00) immediately after the stress-induced sample was taken and then taking a final blood sample (< 30 μl) 30 minutes after injection (Taff et al. 2018). Dexamethasone is a synthetic glucocorticoid that primarily binds peripheral glucocorticoid receptors, stimulating negative feedback. We previously validated this method and dose in tree swallows from this population (Taff et al. 2018; Zimmer et al. 2019). At the third capture, we took only a single blood sample to measure baseline corticosterone and females were released within five minutes. Blood samples were stored in coolers in the field; red blood cells and plasma were separated by centrifugation within 3 hours and then stored at −30° C until processing.

### Plumage Manipulation

At the first capture, females were assigned to either a breast plumage dulling treatment group or a sham control treatment group. These assignments were made randomly except that we balanced the treatments by female age (first time breeders vs. returning breeders) by randomizing treatment and control assignments within each age class. In total, the dulled group included 34 females (18 SY and 16 ASY) and the control group included 36 females (21 SY and 15 ASY). At the first capture, we collected 6-8 breast feathers to measure breast brightness prior to any treatment. For the females in the dulling treatment, we then uniformly colored the breast from just below the beak down to the legs using a light grey non-toxic marker (Faber-Castell PITT artist pen ‘big brush’ warm grey III 272). Test feathers colored with this marker had uniformly lower reflectance across the entire visible spectrum. For females in the control treatment, we applied a colorless non-toxic marker over the same plumage area for the same length of time (Prismacolor premier colorless blender PB-121). For both groups, we collected an additional 6-8 feathers after coloring to assess the immediate impact of each treatment on breast coloration.

Based on previous experiments in other species that used similar methods, we expected the color that we applied to fade over time. Therefore, we collected an additional 6-8 feathers at the 2^nd^ and 3^rd^ captures so that we could assess the duration of color manipulations using this method. After collecting feathers at each of those capture, we also re-applied the dull or clear marker to maximize the amount of time during the breeding season that our color manipulations would be in effect. Thus, each individual was colored a total of three times during the experiment. With the capture schedule that we employed, we were able to assess fading over a 10-14 day period (1^st^ to 2^nd^ captures) and over a 3-5 day period (2^nd^ to 3^rd^ captures).

### Radio Frequency Identification Sensor Network

We used RFID readers to monitor activity at nest boxes to determine both feeding rates and social interaction patterns. Each RFID unit consisted of a 12-volt battery placed on the ground next to the nest box, an RFID circuit board, and a circular antenna that was attached around the outside of the nest box entrance hole. The RFID board and cables connecting to the antenna and battery were housed in a small plastic container that was attached to the bottom of each nest box. The board was programmed to poll for a PIT tag within the antenna radius (a few centimeters from the entrance hole) every second from 5 am until 10 pm each day. The placement of the antenna meant that only individuals that either passed through the entrance hole or perched on the entrance hole would have been recorded. We changed the battery that ran the system every 5-7 days and downloaded data from each box 1-2 times during the season and one final time when units were uninstalled.

We installed RFID readers at each active box on day 4 after clutch completion (2 days before the first capture) so that complete data were available for most nests from day 6 of incubation (when PIT tags were applied) through fledging. Occasionally, RFID boards malfunctioned or batteries failed so that the total number of days varied between boxes and we accounted for these sampling differences in our analyses (see below). At three boxes, board malfunctions or incorrectly saved files meant that we had no RFID data and these nests are excluded from most analyses that use RFID data (though these focal females could still be recorded making trips to other nests). One additional female was excluded from some RFID based analyses because she only had one leg and thus could not be equipped with a PIT tag (this female was still included in analyses looking at visitors to her box).

### Plumage Measurements

We measured the reflectance characteristics of feathers collected from the breast using an Ocean Optics FLAME-S-UV-VIS spectrophotometer with PX-2 pulsed Xenon light source and WS-1 white standard in OceanView version 1.5.2 (Ocean Optics, Dunedin, FL). For each measurement, we stacked and taped four feathers on an index card and then smoothed the barbs to create a patch large enough for measurement. We used a fiber optic UV/VIS probe in a holster that blocked external light and maintained a distance of 5 mm between the feathers and probe. Spectra were collected with a 10 scan average, 20 nm boxcar width, and 60 ms integration time. Four separate spectra were taken for each feather stack with the probe removed between measurements. For each female, we measured four sets of feathers (two from 1^st^ capture and one each from 2^nd^ and 3^rd^ capture; see above).

Reflectance spectra generated by OceanView were processed in R version 3.3.3 (R Core Development Team, 2016) using the package ‘pavo’ (Maia et al. 2013). Based on a prior study (Taff et al. 2019b), we were mainly interested in the overall brightness of the breast and our manipulation was directly relevant for this plumage metric. Thus, we did not consider all the color metrics explored by Taff et al. (2019b) and we instead focused exclusively on breast brightness. We calculated mean breast brightness as the average reflectance from 300-700 nm (‘B2’ in the ‘pavo’ package). The four repeated measurements from each feather sample were averaged to arrive at a final brightness measure.

### Feeding Rate

We used the records from RFID readers installed at each box to record feeding rates for each female and male equipped with a PIT tag in this study (as in Vitousek et al. 2018b). For this analysis, we only included RFID records from days after hatching up until fledging. The readers that we used were programmed to record the presence of a PIT tag every second. Thus, a single feeding trip often produced multiple recordings in close succession and we needed to apply a time threshold to determine feeding rates (i.e., how far apart must two readings be in order to be considered as separate feeding trips).

We followed the process developed by Vitousek et al. (2018b) to determine feeding rates. That study paired video recordings and RFID records to determine the optimal time threshold empirically. Because parental behavior changes as nestlings age (e.g. the amount of time spent perched on the entrance hole changes dramatically), the time threshold determined by Vitousek et al. (2018b) also differs with nestling age (nestling ages 1-3 days: 136.5 seconds; 4-6 days: 55.5 seconds; 7-9 days: 36.5 seconds; 10-12 days: 20 seconds; 13-15 days: 11 seconds; 16-18 days: 25.5 seconds). After filtering the raw RFID records using these time thresholds, we calculated the total number of daily feeding trips for each female and male in the population. Note that males were not captured until day 6 after hatching and some males were never captured. Therefore, feeding data was less complete for males than for females.

### Visitation Patterns

We also used RFID records to derive nest box visitation patterns for each female in our study. To process RFID files, we used the same custom R script that was developed and described in detail in (Taff et al. 2019b). Briefly, this script reads a batch of RFID files, merges and integrates metadata stripped from the file names, and performs a number of data cleaning operations to correct PIT tag reading errors. Once these initial steps are complete, the fully merged file of RFID records is used to determine each instance of a non-box owner visiting another box in the population. As with the feeding rate data above, visits to other boxes often produce a series of readings rather than a single read and these should be considered as a single ‘visit’. We considered readings of the same individual that occurred <120 seconds apart to be part of the same visit. The 120-second threshold was somewhat arbitrary, but Taff et al. (2019b) previously found that time thresholds of 30-600 seconds resulted in qualitatively similar overall patterns of visitation. Analyses that focused on the total number of unique visitors were unaffected by the time threshold choice.

The end result of this script is a table of specific instances when a tagged individual was recorded at a nest box in the study site at which they were not a part of the breeding pair. For clarity, when discussing these instances, we refer to readings from the perspective of a focal breeding female as ‘visits’ when a non-owner visited the box of the focal or ‘trips’ when the focal female made trips to other boxes in the population. For each of these records, we extracted information on the time and duration of the visit as well as the sex, treatment group, and breeding stage of both the visiting and receiving female. The rate of visitation to other boxes differs dramatically with breeding stage (Taff et al. 2019b); therefore, we also used the full visitation table to calculate per-day values for the total number of visits and number of unique visitors of each sex at each box for each day in the breeding season (as in Taff et al. 2019b). In addition to the 70 boxes included in our main study, some additional nesting attempts occurred at our study sites that were not entered into treatment groups and some adults were equipped with PIT tags that were not directly part of this study. For the purposes of our analysis of box visitation patterns, we included interactions with adults from these additional boxes because they provided a more complete picture of the overall social structure of the population.

### Physiological Measurements

We measured pre-and post-treatment values for glucose (baseline and stress-induced) and corticosterone (baseline, stress-induced, and post-dexamethasone). Glucose was measured in the field immediately after baseline and stress-induced blood samples were collected using a FreeStyle Lite blood glucose meter and test strips (Abbott Diabetes Care, Alameda, CA, USA). These devices have been recommended for use in birds (Breuner et al. 2013) and similar models have been used in prior studies of wild birds (e.g., Clinchy et al. 2004; Malisch et al. 2018). In 2018, we assessed the reliability of glucose measurements obtained with this technique by measuring glucose with two independent test strips from the same blood sample for 21 females. Based on those samples, the CV for repeated measurements was 4.4% and repeatability was 0.73.

We measured corticosterone concentration in baseline, stress-induced, and post-dexamethasone plasma samples using commercially available enzyme immunoassay (EIA) kits (DetectX Corticosterone, Arbor Assays: K014-H5). We previously validated these kits in tree swallows and extensive validation and protocol details are available in Taff et al. (2019b). Briefly, we used 5 μl of plasma in a triple ethyl acetate extraction and then ran the resulting samples in duplicate following the manufacturer’s protocol. Extraction efficiency was determined using samples spiked with a known amount of corticosterone; average extraction efficiency with this method was 89.7%. When starting with 5 μl of plasma, the lower detection limit was 0.8 ng/μl. Inter-plate variation was assessed using a plasma pool run across plates and was 5.7%. Intra-plate variation was assessed using duplicate wells and averaged 10.6%.

### Microbiome Sample Processing

We extracted DNA from cloacal swabs using DNeasy PowerSoil DNA Isolation Kits (Qiagen Incorporated) following the manufacturer’s protocol. To amplify the 16s rRNA gene we closely followed the Earth Microbiome Project 16s Illumina Amplicon protocol (Caporaso et al. 2012; Caporaso et al. 2011), with the exception of using 10 μl total reaction volumes instead of 25 μl. We amplified the V4 region of the 16S gene using the primers 515F and 806R with Illumina adapters added. Each PCR reaction was run in triplicate and included 5 μl of 2x Platinum Hot Start Master Mix (Invitrogen), 0.5 μl of 10 μM primers, 3 μl of nuclease free water, and 1 μl of template DNA. Cycling conditions were 3 minutes at 94° C followed by 35 cycles of 94° C for 45 seconds, 50° C for 60 s, and 72° C for 90 seconds before a final extension at 72° C for 10 minutes.

The three replicate reactions for each sample were pooled and run on a 1% agarose gel to confirm that amplification was successful. Each PCR run included negative controls (nuclease free water in place of template DNA). We also extracted and amplified 5 negative control swabs. For these negative controls we added a sterile swab directly to RNAlater during the field season and then stored and processed the swabs exactly as described above for true samples. We submitted our final pooled PCR products to the Cornell Biotechnology Resource Center for quantification, normalization, library preparation, and sequencing. Our samples and negative controls were sequenced in one Illumina MiSeq PE 2 x 250 bp run.

### Microbiome Bioinformatics

We processed raw sequence data from microbiome samples in R following the microbiome workflow described by Callahan et al. (2017). Briefly, after visually inspecting quality scores for our sequences, we removed forward and reverse primers and truncated reads at 180 bp using the default settings in the ‘filterAndTrim’ function of the ‘dada2’ package in R (Callahan et al. 2016). We proceeded through the standard workflow in ‘dada2’ including dereplication, modeling sequencing error based on our data, calling amplicon sequence variants (ASVs), merging forward and reverse reads, and removing chimeras using default settings to arrive at a final set of identified ASVs for each sample. From this ASV table, we made taxonomic assignments based on the Silva 132 database (Quast et al. 2013; Yilmaz et al. 2014). We next built a generalized time-reversible maximum likelihood phylogenetic tree with Gamma rate variation using the ‘phangorn’ package in R (Schliep 2010). We combined the ASV table, sample data, taxonomy table, and phylogenetic tree for subsequent analysis using the ‘phyloseq’ package (McMurdie & Holmes 2013). Finally, after removing eukaryotes, archaea, chloroplasts, singletons, and mitochondria, we used the ‘decontam’ package in R to identify and remove likely contaminants based on associations with sample biomass and comparisons with negative controls using the default settings (Davis et al. 2018).

From this final ASV file we determined the relative abundance of different taxa agglomerated at the phylum and family level. We compared the composition of microbiome samples from control and dulled birds using nonmetric multidimensional scaling based on weighted unifrac distances and permanova tests using the ‘adonis’ function from the ‘vegan’ package in R. We also calculated three alpha diversity metrics using the ‘phyloseq’ and ‘picante’ packages in R: the Shannon Diversity Index, Faith’s Phylogenetic Diversity (PD), and Simpson’s Diversity Index (Kembel et al. 2010; McMurdie & Holmes 2013). To calculate alpha diversity we had to rarify our samples to an even depth of sequencing coverage. Although most samples had ample coverage (mean reads per sample = 41,325 ± 26,421), some samples did return a relatively small number of reads (e.g., samples with < 1,000 reads = 11, < 500 = 6, < 250 = 3).

Because we planned to include pre-and post-treatment measures in our models we did not want to set an arbitrarily high rarefaction limit and risk losing a substantial part of our sample. Thus, we initially calculated diversity metrics using a series of rarefaction limits ranging from the minimum number of sample reads (153) up to 20,000 reads using the ‘prune_samples’ function in ‘phyloseq’. Although the total number of ASVs detected increased with higher read depth, the variation in diversity metrics derived from only 153 samples was very similar to that obtained from 20,000 samples (pairwise correlation coefficient comparing 153 vs. 20,000 read rarefaction limit: Shannon = 0.99, Simpson = 0.99, PD = 0.72). Thus, we rarefied at this lower limit to allow for inclusion of all of the experimental birds in our study. We repeated this analysis with a rarefaction limit of 1,500 and despite the smaller sample size the overall results were qualitatively similar. For simplicity, we present results only using the lower rarefaction limit.

### Data Analysis

In our main analysis, we were interested in identifying the total causal effects (if any) of breast coloration on conspecific behavior, focal individual behavior and physiology, and, ultimately, seasonal reproductive success. We assumed that females could not directly assess their own plumage; therefore all causal pathways from plumage to performance are indirect and necessarily pass through receiver behavior. Because we directly manipulated plumage, we interpret any effect of dulling as a causal result of plumage expression, even if the exact indirect path is unknown (Pearl 2009). Thus, we initially fit a series of models for each measured response variable that included only the main factors of interest: treatment (control vs. dulled), initial brightness, and a treatment by brightness interaction. We included the treatment by brightness interaction because we expected that initially bright females would experience a greater magnitude change in coloration after dulling compared to initially dull females and the consequences of dulling might therefore differ. Based on the results obtained in these initial models, we next explored the evidence supporting several plausible indirect pathways connecting plumage expression and performance.

While the main predictor variables included in the initial models for each response variable were similar, the structure of the models differed for some response variables based on the type of data. For behavioral data collected from the RFID network, we had many days of observations for each individual nest. We were interested in fixed effects (treatment, initial brightness) at the level of individuals, but expected covariates at the level of observations (day in breeding cycle, date in season) to have a large effect on individual observations. Thus, we fit these responses as generalized linear mixed models (GLMMs) or linear mixed models (LMMs) that included our predictors of interest but also accounted for observation level differences. Specifically, we added a random effect for day of the breeding cycle (to account for breeding stage related changes in behavior) and a random effect of date in the season (to account for variation in behavior associated with daily conditions such as temperature or food availability) in each model. We were not interested in directly interpreting these random effects and used them only to account for behavioral changes that are known to occur through the breeding cycle.

Using the same model structure described above, we tested a total of ten behavioral response variables. First, we fit four models that described conspecific visitation behavior as the total number of visits and number of unique visits to the focal box on each day of observation; this metric was calculated separately for male and female visitors, both because the function of signals might differ between the sexes and because PIT tags were not deployed on as many males.

Next, we fit two models with male provisioning rate (number of feeding trips on each day) and total time spent at the nest entrance (raw number of reads) as the response variables. We then fit four models describing aspects of the focal females behavior: provisioning rate, time spent at the nest entrance during incubation, and number of unique or total trips to other boxes in the population on each day of observation. Provisioning rate (for males and females) and time at the entrance were fit as LMMs and focal female trips to other boxes were fit as GLMMs with a Poisson error distribution. We interpreted the effect of our predictors from the fully specified models, but for plotting purposes we also extracted best linear unbiased predictors (BLUPs) from these models to allow for plotting of behavior versus treatment and plumage measurements.

We initially fit four GLMMs with the total number of visits or number of unique visitors by males or females as the response variable with a Poisson error distribution using the ‘glmer’ function in the ‘lme4’ package (Bates et al. 2015). These models included a treatment by initial brightness interaction as main effects along with day in the breeding stage and date as random effects. These specifications were retained for the models of unique visitors, but models for the total number of visitors were overdispersed and zero-inflated as a result of some days with many repeated visits by the same individuals. Thus, we refit those responses as zero-inflated negative binomial models using the ‘glmmTMB’ package (Brooks et al. 2017). All models also included an individual identity random effect to account for repeated observations and an observation level random effect to correct for overdispersion.

For female physiological measures and microbiome diversity, we fit a set of four models with post-treatment measures as the response and pre-treatment measures, treatment, and a treatment by initial breast brightness interaction as predictors. Finally, for females that returned to breed in the year after manipulations, we compared the date of nest initiation and clutch size in year *n + 1* to treatment group in year *n* using a simple linear model. We did not compare other reproductive measures in year *n + 1* because most females that returned were part of a separate experiment in that year that started after eggs were laid.

In all of our analyses, we considered effects with 95% confidence intervals that do not overlap zero to be meaningful. Whenever possible, we standardized predictor and response variables to a mean of 0 and standard deviation of 1 so that effect sizes are in units of standard deviations. All analyses and figures were produced in R version 3.5.1 (R Core Development Team).

## RESULTS

### Plumage Dulling Treatment

The plumage dulling treatment reduced overall brightness of breast feathers without altering other spectral characteristics (Figure S1). We compared pre and post treatment brightness for control and dulled females using a single LMM with brightness as the response variable. The model included fixed effects for treatment, capture number, and their interaction as well as a random effect for female identity. From the fit model, we determined the difference in brightness (in units of standard deviations) between initial coloration and samples collected 1) immediately post treatment, 2) 10-14 days post treatment, and 3) 3-5 days post treatment for both control and dulled females.

At the initial capture, females assigned to control and dulling groups did not differ in pretreatment brightness (control - dulled; mean = −0.01, CI of difference = −0.47 to 0.44). Control female brightness did not differ from initial brightness in any subsequent samples (initial vs. immediate post: mean of difference = −0.03, CI of difference = 0.42 to −0.37; initial vs. 10-14 days: mean = −0.25, CI = 0.66 to −0.16; initial vs. 3-5 days: mean = −0.25, CI = 0.68 to −0.17). In contrast, dulled female brightness was lower than initial brightness at every post-treatment time point, though the difference declined with increasing time between sampling as expected if the color faded after application (initial vs. immediate post: mean = −0.99, CI = −0.58 to −1.39; initial vs. 10-14 days: mean = −0.49, CI = −0.08 to −0.90; initial vs. 3-5 days: mean = −0.64, CI = −0.21 to −1.08).

### Conspecific Social Interactions

Our RFID system recorded social activity from 8 days pre-hatching (just after treatments were initially applied) to 16 days post-hatch. In total, we identified 18,919 instances in which a known bird was recorded at a box where they were not part of the breeding pair. Both males and females in the population regularly visited boxes included in our study at which they were not breeding (males: 11,297 visits; females 7,622).

The best-supported model for the total number of visits by conspecific males per day included a three-way interaction between treatment, initial plumage brightness, and breeding stage (Table S1; model weight = 0.70). Focal females that received the dulling treatment received fewer total male visitors, but the effect was only apparent among females that were initially bright and was more pronounced at later breeding stages (Figure 1 A & B; Table S2). For unique male visitors, two models received equal support; these models included either the full three-way interaction, or twoway interactions with treatment and both initial brightness and breeding stage (Table S1; model weight for full model = 0.37, two-way interaction model = 0.38). In this case, focal females received fewer unique male visitors after the dulling treatment, but only if they were initially bright; the uncertainty in model selection results from uncertainty in whether the effect of this interaction differed across breeding stages (Table S2; Figure S2). For both total and unique male visitors, the rate of visitation tended to be higher at later breeding stages and among brighter females, independent of the dulling treatment.

**Figure 1.**
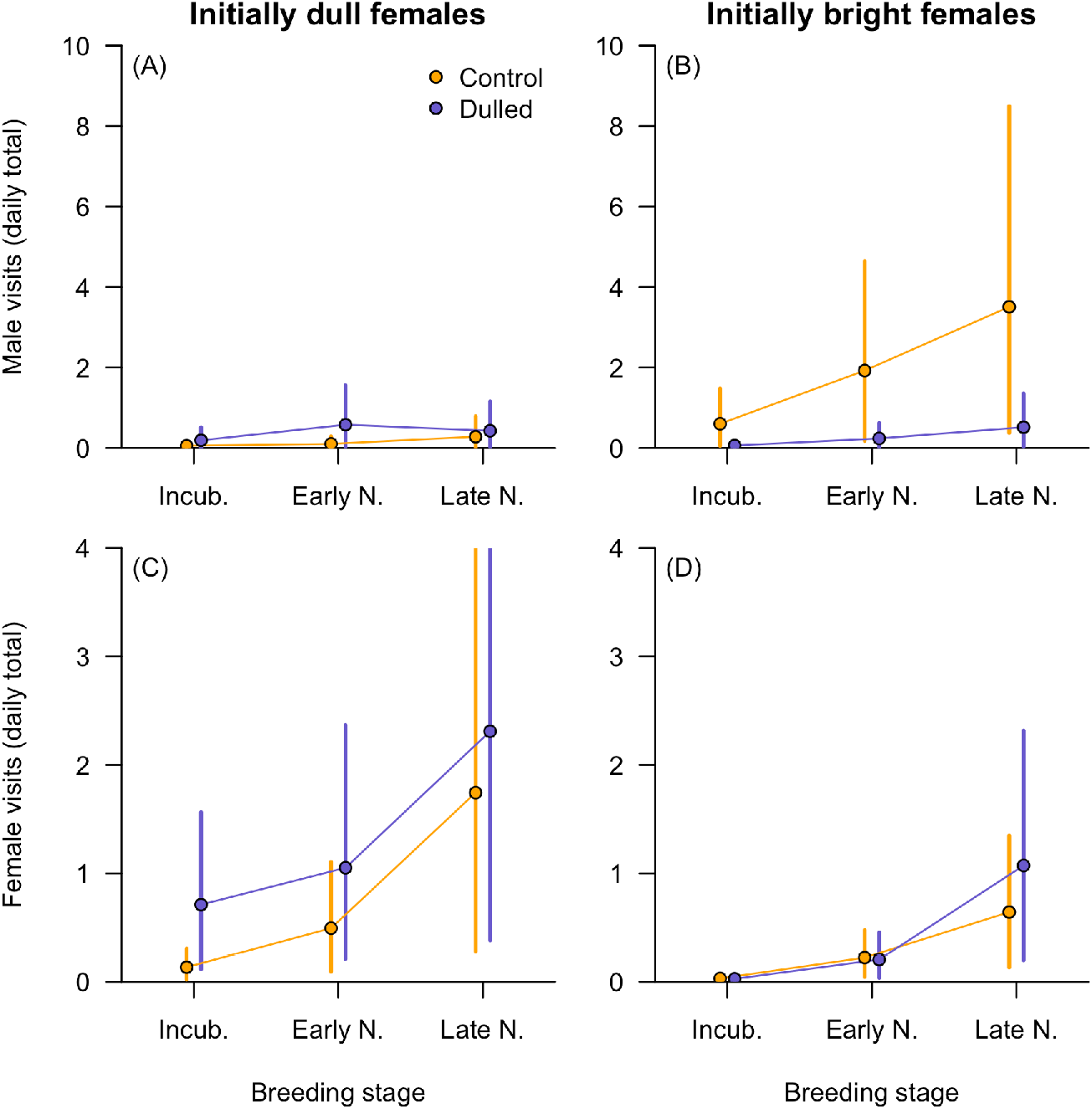
Model predicted illustration of the three-way interaction between treatment, breeding stage (incubation, early nestlings, and late nestlings), and plumage treatment based on the best-supported models for total number of male visitors (A & B) and total number of female visitors (C & D). The paired panels illustrate predictions for females that were initially 1 SD below the mean (A & C) or 1 SD above the mean (B & D) for breast brightness. Male visit rate did not differ between treatment groups when females were initially dull (A), but was higher for control females that were initially bright and the effect was more pronounced at later breeding stages (B). Female visitation rate was higher when initially dull birds received a dulling treatment and this effect was most pronounced during incubation (C); for initially bright females, female visitation rate did not differ with treatment but did increase across breeding stages (D).

Among candidate models for the total number of daily visits by conspecific females, the full model that included a three-way interaction between treatment, initial plumage brightness, and breeding stage received strong support (Table S1; model weight = 0.99). The same model specification received the most support for the number of unique female visitors per day (Table S1; model weight > 0.99). In both cases, focal boxes received more visits by females (or more unique visitors) when the focal female was initially dull and visit rate increased as breeding stage progressed. Focal females that received the dulling treatment tended to receive more female visitors if the focal female was initially dull and this effect was most pronounced during incubation (Figure 1 C & D, Figure S2, Table S2).

### Behavior of the Social Mate: Time at Entrance and Provisioning

We restricted our analysis of the behavior of experimental females’ social mates to the 7 to 16 days after hatching because most males did not have PIT tags before that time and because that is the period where most nestling provisioning occurs. After applying these criteria, we recorded a total of 302 days of activity from 41 different males that included 191,856 RFID records and 51,969 identified feeding trips at the focal box. We fit two candidate sets of LMMs with total time at the entrance hole (raw number of reads each day) or feeding rate (daily feeding trips) as the standardized response variable. Fixed effects included focal female initial brightness, treatment, and their interaction. All models except the intercept only model also included nestling age and a quadratic effect of nestling age because feeding rate is known to increase steadily as nestlings age and then decline again as nestlings approach fledging age (Vitousek et al. 2018b). Each model included random effects for nest identity and date in the breeding season.

The best-supported model for total male time at the nest entrance included only nestling age and the quadratic effect of nestling age (Table S3, model weight = 0.85). The same model also received the most support for male provisioning rate (Table S3, model weight = 0.77). Thus, while males increased their activity and feeding rate around the nest box as nestlings aged, there was no evidence that either of these behavioral measures were influenced by the treatment that their mates received.

### Focal Female Behavior: Time at Entrance, Provisioning, and Interactivity

For focal females, we recorded time at the nest entrance (raw number of reads each day) and feeding rate (daily feeding trips) throughout the reproductive period. We restricted models to observations made from the day after hatching through day 16 after hatching. Applying these criteria, we recorded a total of 752 days of activity from 59 females that included 490,536 RFID records and 124,281 identified feeding trips.

We found no evidence that female time at the nest entrance differed with respect to treatment. The best-supported model for the total number of daily female RFID reads included only the intercept (Table S4, model weight = 1). For provisioning, the best-supported model included treatment along with nestling age and the quadratic effect of nestling age (Table S4, model weight = 0.76). All females increased their provisioning rate as nestlings aged, but experimentally dulled females consistently fed nestlings at a higher rate (Figure 2; Table S5; dulling β = 0.35, CI = 0.09 to 0.60)

**Figure 2.**
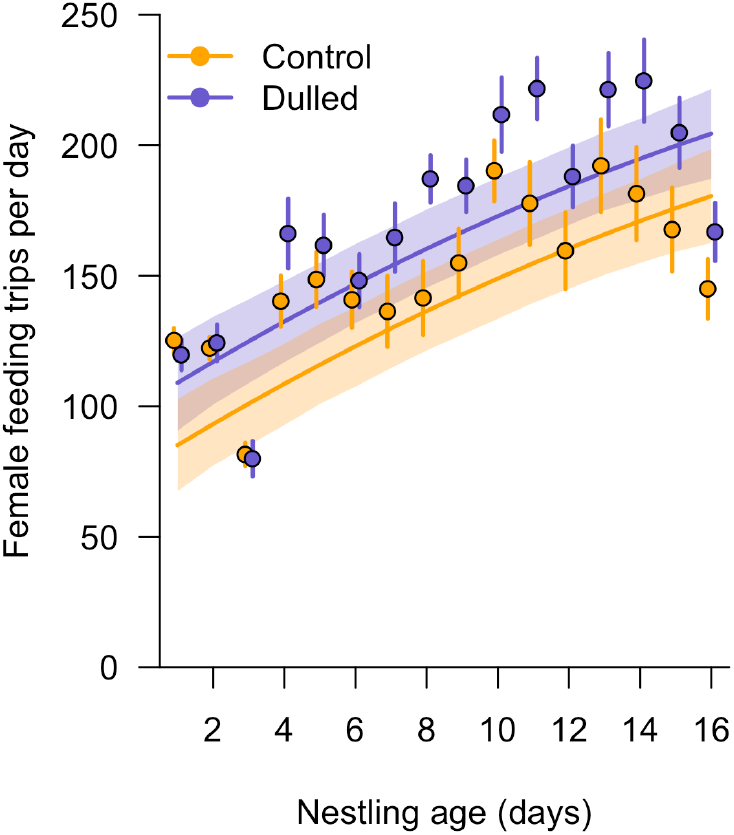
Relationship between nestling age and female provisioning rate for control and dulled females. Points and vertical lines are the mean ± SEM of the raw data at each nestling age. Shaded regions and trend lines are model predicted relationships based on the best-supported model in Table S4.

We fit candidate model sets of GLMMs for the total number of trips made and number of unique trips made each day for focal females (total trips as zero-inflated negative binomials and unique trips as Poisson). In 1,223 days of observation, we recorded a total of 7,307 trips made by 64 focal females to other boxes in the population at which they were not breeding (note that this number includes trips to some monitored boxes that were not part of this study). For both the total number of daily trips made and the number of unique trips made each day, the best supported model was the full model that included a three-way interaction between treatment, initial brightness, and breeding stage (Table S6, model weight > 0.99). In both cases, females that received the dulling treatment made fewer total trips to other boxes, but the magnitude of this effect was greater when females were initially dull and at later breeding stages (Figure 3, Figure S3, Table S7).

**Figure 3.**
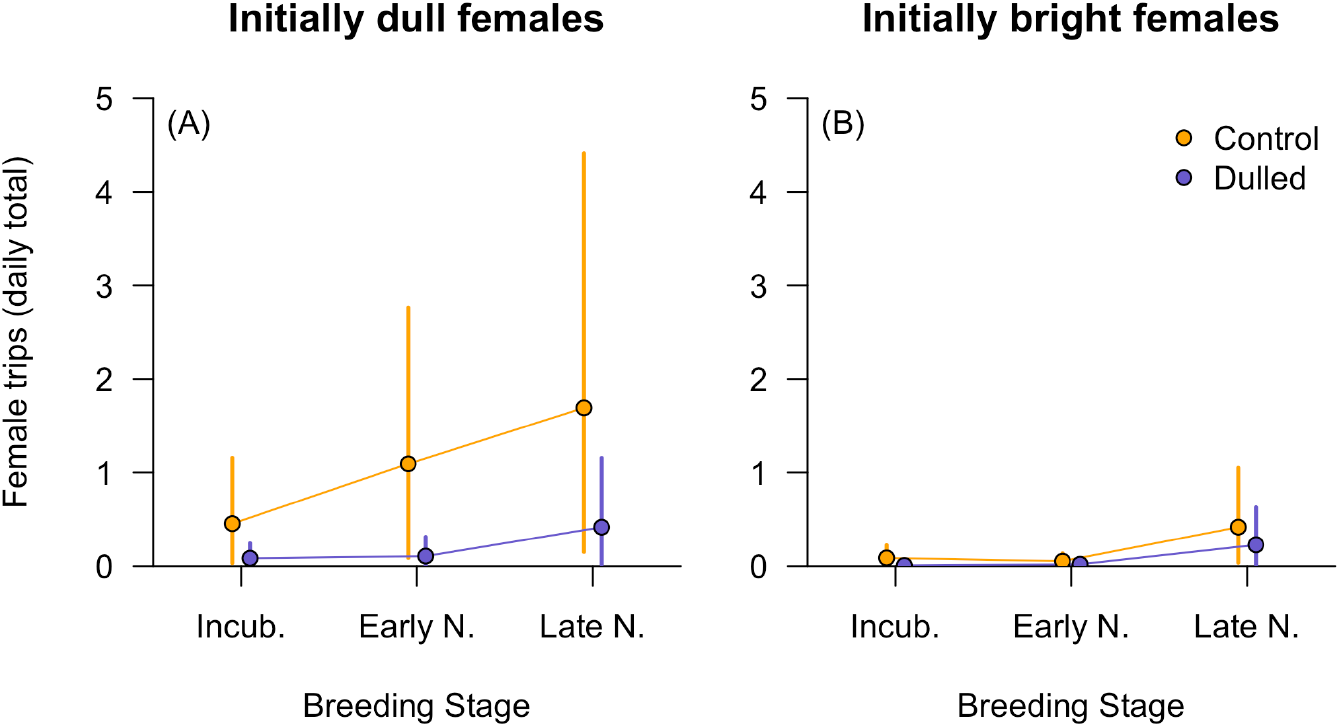
Illustration of the three-way interaction included in the best-supported model for female trips to other boxes. The two panels show the relationship between treatment and number of trips made at different breeding stages for females that were initially 1 SD below the mean for plumage brightness (A) and 1 SD above the mean (B). Females that were dulled made fewer trips to other boxes, but this effect was only evident when females were initially dull and the magnitude of the effect increased at later breeding stages.

### Focal Female Physiology and Condition

For each physiological variable that we measured we fit a linear model that included combinations of the pre-treatment measure, initial plumage brightness, treatment, and the treatment by initial brightness interaction. In total, we fit six models with this same structure and with the response variables of measures taken at the second capture (baseline, stress-induced, and post-dexamethasone corticosterone, baseline and stress-induced glucose, mass) or at the third capture (baseline corticosterone and mass). We found no evidence that dulling predicted post-treatment baseline corticosterone, stress-induced corticosterone, dexamethasone-induced corticosterone, baseline glucose, or mass at any time point (Table S8).

For stress-induced glucose levels at the second capture, the best-supported model included an interaction between initial coloration and plumage treatment (Table S8; model weight = 0.70); in dulled females, stress-induced glucose at this capture was positively related to initial brightness, but there was no relationship in control females (Figure 4; Table S9). This positive relationship was largely driven by the fact that initially dull birds that received the dulling treatment had lower stress-induced glucose levels at the second capture.

**Figure 4.**
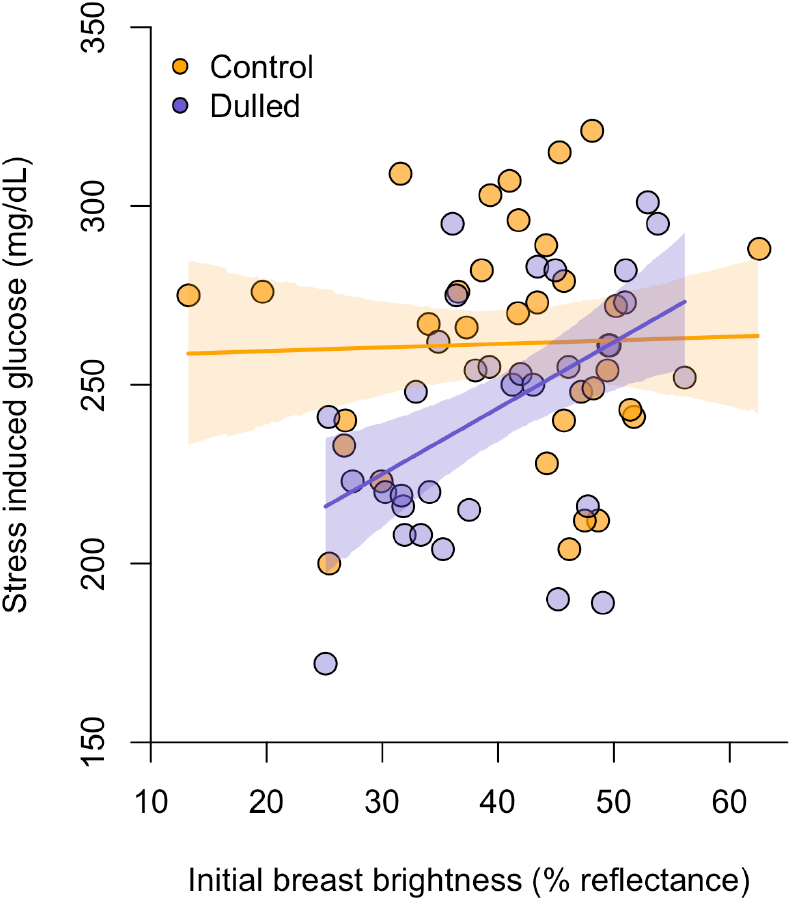
Relationship between pre-treatment breast brightness and post treatment stress-induced glucose for control vs. dulled treatment groups. Points are raw data, but model predicted best fit lines and confidence intervals are based on the best-supported model (Table S9) that controls for pre-treatment measurements.

### Focal Female Microbiome

We fit a set of candidate models for microbiome alpha-diversity (Shannon, Simpson, and PD) at the second and third capture. For Shannon and Simpson diversity, the best-supported model at the second capture included only the intercept. For PD, the best-supported model at the second capture included treatment and initial brightness, but the intercept only model received a similar level of support (Table S10). However, at the third capture, the best-supported model for all three metrics included pre-treatment diversity and treatment (Table S10; model weight for Simpson diversity = 0.78; Shannon diversity = 0.71; PD = 0.77). For all metrics, females that received the dulling treatment had higher microbiome diversity at the third capture (Figure 5A; Table S11; Simpson dulled β = 0.70, CI = 0.17 to 1.22; Shannon dulled β = 0.70, CI = 0.16 to 1.24; PD dulled β = 0.32 to 1.36).

**Figure 5.**
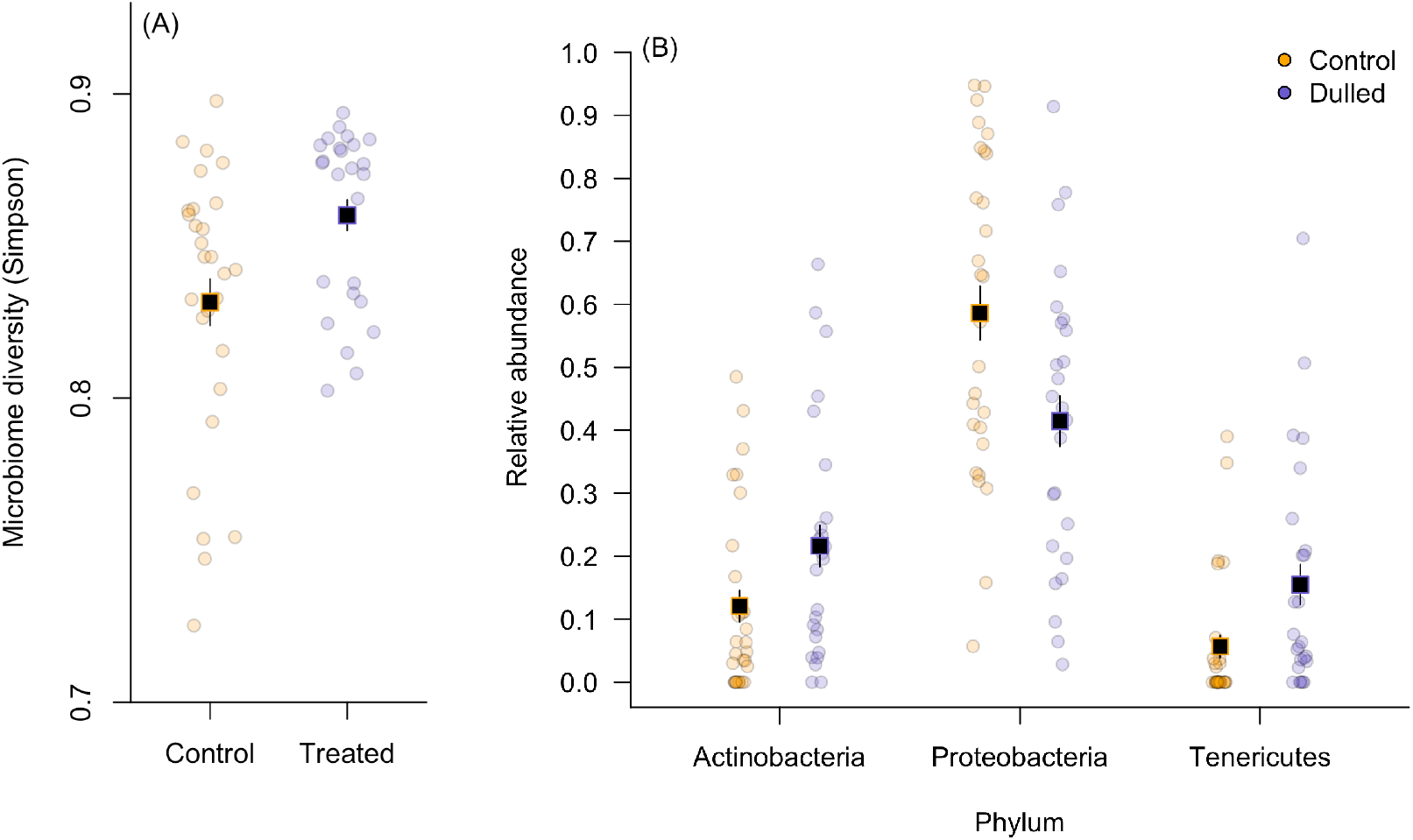
(A) Simpson diversity at the third capture for females that received the dulling or control treatment. (B) Relative abundance of three common phyla in control vs. dulled females at the third capture. In both panels, points illustrate raw data while vertical lines and squares represent estimates and 95% confidence intervals based on the best-supported models in tables S10 and S12.

When comparing weighted unifrac distances, females in the control and dulled treatment groups did not differ at the first and second capture (Figure S4; first capture PERMANOVA F_1,62_ = 19.5, P = 0.15; second capture F_1,62_ = 0.27, P = 0.93). However, at the third capture, there was a significant difference between the microbiome communities measured in control and dulled females (Figure S4; PERMANOVA F_1,51_ = 3.5, P = 0.003).

Overall, the most common phyla that we detected were Proteobacteria (37% of sequences), Actinobacteria (24%), Firmicutes (19%), Tenericutes (16%), and Bacteroidetes (2%; Figure S5). For each of these phyla we fit a similar set of candidate models to those described above. The best-supported model for relative abundance of Firmicutes included only pre-treatment abundance (Table S12), but the best-supported model for the other four common phyla included treatment (Table S12). By their third capture, females that received the dulling treatment had lower relative abundance of Proteobacteria (Figure 5B; Table S13; dulled β = −0.16, CI = −0.29 to −0.02) and higher relative abundance of Actinobacteria, Bacteroidetes, and Tenericutes (Figure 5B; Table S13; Actinobacteria dulled β = 0.10, CI = 0.0 to 0.19; Bacteroidetes β = 0.02, CI = 0.0 to 0.04; Tenericutes β = 0.08, CI = 0.01 to 0.15). While the model weights and confidence intervals for these effects indicate some uncertainty, the estimated magnitude of the effects was relatively large, as the three rarer phyla were about twice as abundant in dulled compared to control females by the third capture. An analysis of relative abundance at the family level yielded qualitatively similar results (Figures S6-8).

### Focal Female Reproductive Success and Nestling Phenotype

Overall, females in the dulled treatment group fledged more offspring and had more offspring in the nest at each reproductive stage (Figure 6). We fit a set of three binomial GLMMs with individual nestling fate as the binomial response, nest identity as a random effect, and treatment, initial brightness, or their interaction as predictors. We fit this set of models separately for survival to hatching, day 12, or fledging to estimate the total effect of treatment on reproductive performance at each of these stages.

**Figure 6.**
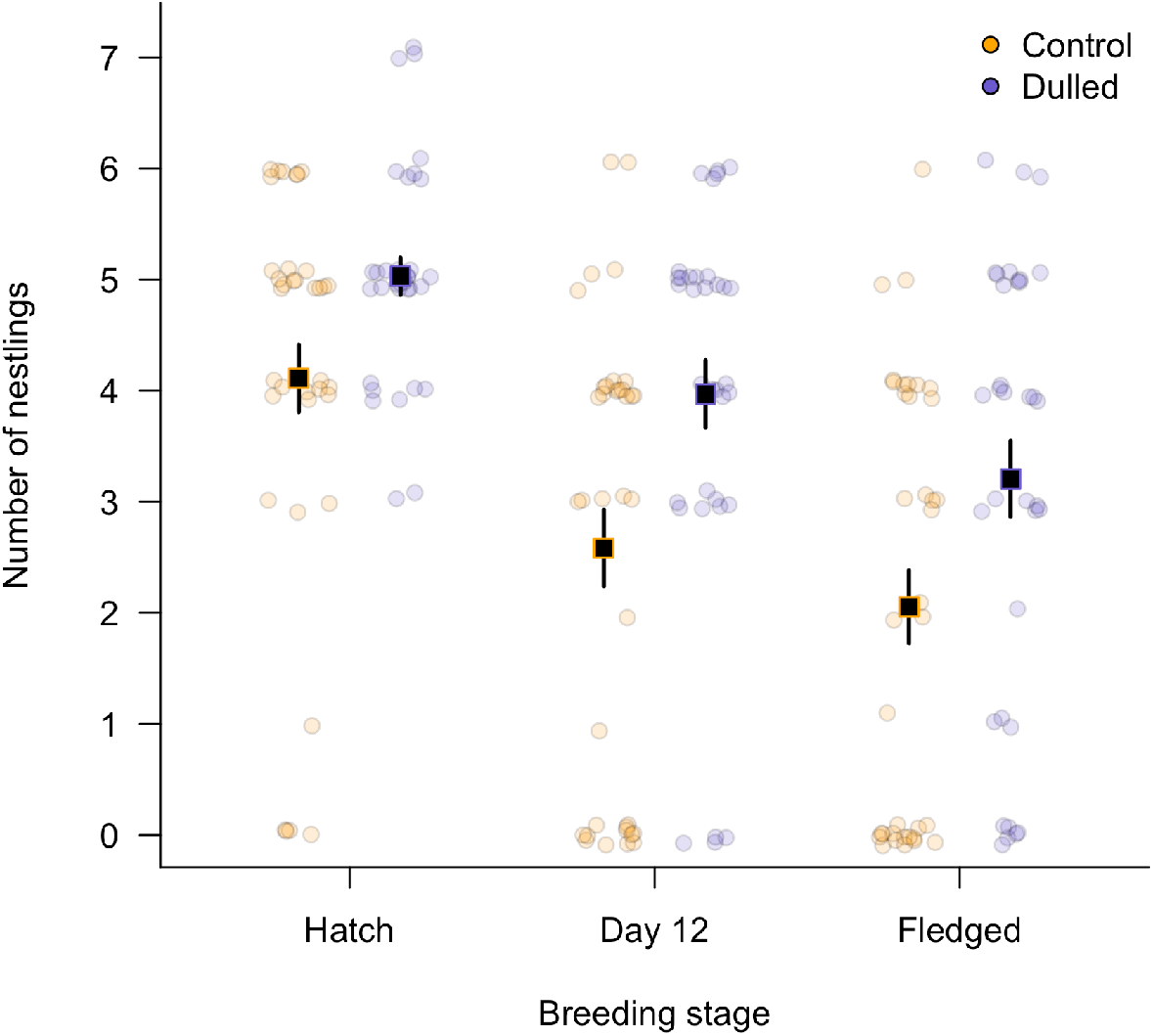
Number of nestlings alive at each time point in nests that received control or dulling treatments. Circles are raw data jittered for clarity, squares are means ± 1 SEM.

For all response variables, the best-supported model included a main effect of treatment, though there was uncertainty over the inclusion of an interaction between treatment and initial brightness (Table S14; combined model weight for top two models that include treatment for hatching success = 0.89; survival to day 12 = 0.96; survival to fledging = 0.81). In all cases, the confidence interval for the interaction effect and main effect of initial brightness crossed zero (Table S15). In the best-supported model for hatching success and survival to day 12, the dulling treatment was consistently associated with increased survival (Table S15; treatment hatching success β = 1.48, CI = 0.16 to 2.79; survival to day 12 β = 1.78, CI = 0.41 to 3.14). A similar relationship was observed for survival to fledging, but in this case the confidence interval crossed zero, indicating that the survival advantage for nestlings in the dulling treatment was primarily the result of enhanced survival earlier in nestling development (Table S15; survival to fledge β = 1.34, CI = − 0.10 to 2.78).

We found no evidence that the phenotype of nestlings that survived to be sampled on day 12 differed by treatment group. For nestling day 12 mass, wing length, and head + bill length, the best-supported model included only the intercept (Table S14; model weight > 0.7).

### Relationships Between Downstream Effects of Treatment

Based on the total causal effects of dulling treatment identified above, we subsequently asked whether the treatment differences that we observed in feeding rate, glucose metabolism, and microbiome diversity were also directly related to nestling characteristics. Since we did not manipulate these downstream measures directly, we cannot determine causality because multiple causal paths could generate associations. Rather, we fit these models in an effort to understand whether the differences in behavior and physiology represent plausible causal mechanisms to explain the differences in reproductive success that were observed as a result of plumage dulling. Because these analyses were more exploratory, we fit simple models with nestling characteristics (number of nestlings alive on day 12 and at fledging; nestling mass, head + bill, and wing length) as the response and female feeding rate, glucose response, or microbiome diversity as a single predictor.

For feeding rate, we used the best linear unbiased predictor (BLUP) of female feeding rate from the best-supported model above (Table S4) as a single measure of overall differences in feeding rate controlling for nestling age. In simple LMMs with nest identity as a random effect, female feeding rate was not associated with nestling mass, wing, or head + bill measurements on day 12 (Table S16). However, females that had higher feeding rates had more nestlings alive on day 12 and fledged more nestlings overall (Figure 7A; Table S16; number on day 12 feeding rate B = 1.49, CI = 0.35 to 2.63; number fledged B = 2.10, CI = 0.94 to 3.26).

**Figure 7.**
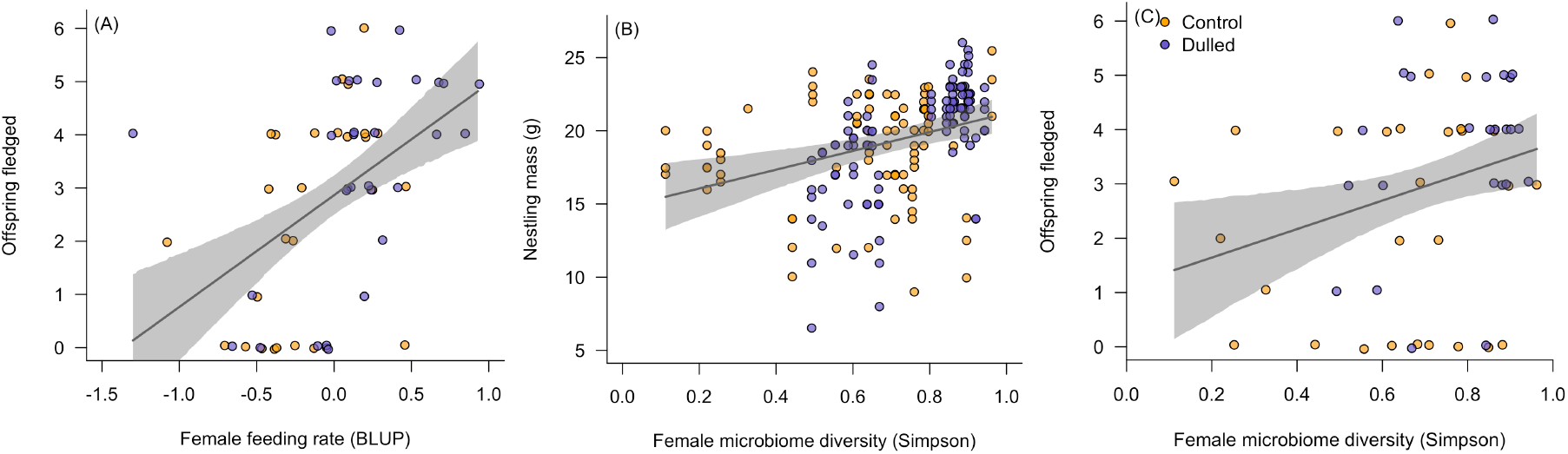
Relationships between (A) female feeding rate and number of offspring fledged, (B) female microbiome diversity and nestling mass on day 12, and (C) female microbiome diversity and number of offspring fledged. Points show raw data and best fit lines and confidence intervals are based on the models in table S16. Points are colored to indicate which treatment females received, but these models did not include treatment directly.

There was a trend for females with higher stress-induced glucose at the third capture to have nestlings that were lighter on day 12, but the confidence interval for this effect spanned zero (Table S16; stress-induced glucose B = − 0.79, CI = −1.68 to 0.11). Stress-induced glucose was unrelated to nestling structural size or to the number of nestlings alive on day 12 or at fledging (Table S16).

Female microbiome diversity at the third capture was positively associated with the mass of nestlings on day 12 (Figure 7B; Simpson B = 6.41, CI = 1.97 to 10.83), but was not related to nestling structural size (Table S16). Microbiome diversity was not related to the number of nestlings alive on day 12, but females with higher microbiome diversity fledged more offspring overall (Figure 7C; Simpson B = 2.62, CI = 0.09 to 5.15).

### Carryover Effects

In total 22 females returned to breed at our site in year *n* + 1 (10 of 34 dulled and 12 of 36 control females). There was no difference in the clutch initiation date in year *n* + 1 for females that had been dulled in the previous year (linear regression with standardized initiation date as response and treatment as categorical predictor: B = −0.27, CI = −1.26 to 0.71). However, females that had been dulled in the previous year laid more eggs in year *n* + 1 (Figure 8; effect of treatment on year *n* + 1 clutch size; B = 0.81, CI = 0.19 to 1.43).

**Figure 8.**
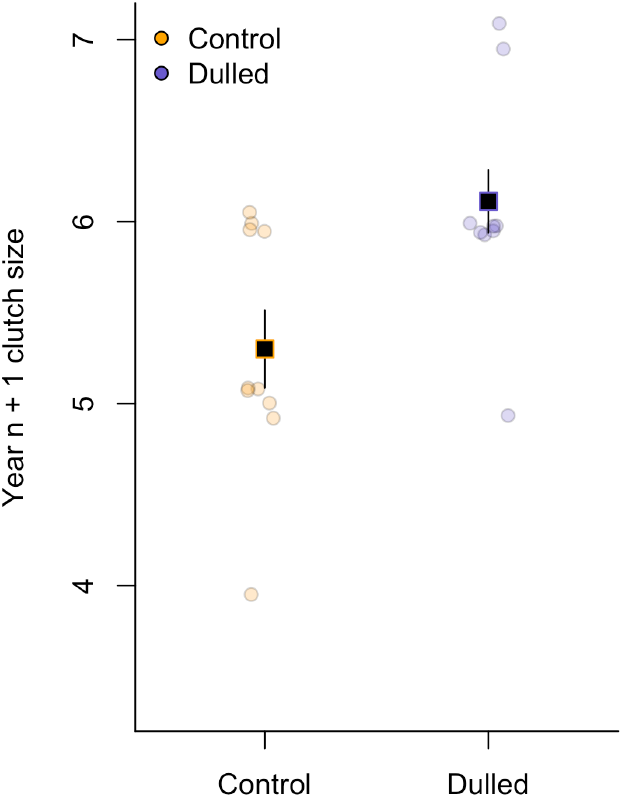
Clutch size in year *n* + 1 for females that received control (orange) or dulled (blue) treatments in year *n*.

## DISCUSSION

We found that experimentally dulling female tree swallow plumage affected a suite of integrated traits. Specifically, females that were dulled experienced fewer male but more female visitors at their own nest, made fewer trips to other nests, fed their nestlings at a higher rate, had a more diverse internal microbiome, and differed in their glucose metabolism relative to sham controls. As a result of these differences, dulled females achieved higher seasonal reproductive success and altered reproductive investment in the subsequent breeding season. Our results demonstrate that signal variation alone—independent of any association with intrinsic differences in condition or genotype—creates feedback between the social environment, physiology, and behavior, resulting in alterations to phenotypic integration. Assuming that females cannot directly assess their own plumage, these patterns must be driven by changes in the experienced social environment that result in females altering their own behavior and physiology. While the exact causal route linking signal expression to reproductive success is not yet known, our results suggest that changes in reproductive investment decisions and internal microbial diversity might play an important role.

One of the major challenges in demonstrating dynamic feedback generated by signals has been that it requires a detailed time series of both physiological and behavioral measurements coupled with signal manipulation (Tibbetts 2014; Vitousek et al. 2014). Our RFID sensor network allowed us to record >18,000 instances in which focal nest boxes were visited by conspecifics. Because tree swallows frequently interact while flying, our recorded social interactions likely represent a subset of those experienced through the entire breeding season. While we could not collect data on the exact nature of these interactions, recent studies in other tree swallow populations have demonstrated that plumage can mediate the frequency of aggressive interactions. Females that have immature plumage are less likely to be the targets of direct aggression (Coady & Dawson 2013) and altering a female’s blue-green back coloration influences nest box retention and reproductive success (Berzins & Dawson 2016, 2018; Berzins et al. 2019). Moreover, females that possess brighter white breast plumage respond more aggressively to a simulated intrusion by another female (Beck & Hopkins 2019). The effects of a single social interaction in this species are most likely small, but the cumulative effects across thousands of interactions throughout a season could be substantial. That dulled females experienced a different social environment than controls indicates that plumage brightness directly mediated conspecific responses to signaling.

In our experiment, females that were dulled achieved higher seasonal reproductive success, but a prior study in the same population found that naturally brighter females were more successful breeders (Taff et al. 2019b). This difference raises the question of why females maintain bright plumage when experimental dulling results in higher performance. One possibility is that the socially mediated costs associated with signal expression are time and context dependent. For example, in the black-crested titmouse (*Baeolophus atricristatus*) the head-crest is a badge of status associated with dominance, but the association is only detectable at times of high competition (Queller & Murphy 2017). We dulled females only after they had begun incubation, but previous work in tree swallows has demonstrated that early season aggression is an important determinant of access to nesting resources (Rosvall 2008, 2011). If signals are important for securing a nest cavity or in early season mate choice then females in our study may have benefited from bright plumage early, but avoided costs of signals that would have accrued later in the season, allowing them to invest more in their own reproductive effort. Variation in the costs and benefits of social signals across time and contexts could contribute to the maintenance of variation in signal expression and to the evolution of different signaling investment strategies.

At this point, we know that plumage manipulation alters the frequency of social interactions, but it is unclear if the nature of those interactions also differs. Our data do suggest, however, that the manipulation had different effects on male and female receivers. Males visited initially dull and experimentally dulled females less often than control females that were initially bright. We do not know if female plumage plays a direct role in mate choice in this species, but tree swallows do have very high levels of extra-pair paternity (Kempenaers et al. 1999; Whittingham & Dunn 2001) and one possibility is that males assess plumage when allocating mating effort. If this is the case, then experimentally dulled females might have escaped continued harassment by males later in the breeding season, allowing them to invest more in reproductive effort. This pattern is similar to that seen in barn swallows (*Hirundo rustica),* where males with experimentally reduced ornamentation increased their provisioning rate (Hasegawa & Arai 2015).

In contrast, conspecific females visited experimentally dulled females more often. This increased visitation rate might have been related to female-female competition, although our manipulations were all conducted after nest settlement. Another possibility is that females were prospecting for public information about neighborhood breeding success. For example, in a study of collared flycatchers (*Ficedula albicollis*), parental activity near the nest box was one of the best predictors of visitation rate by conspecifics (Doligez et al. 2004). In this case, the fact that dulled females were feeding at a higher rate and had more nestlings might have contributed to the increased rate of visits. Future work should focus on complementing our continuous behavioral monitoring with detailed observations of individual social interactions.

Our experiment adds to recent studies that have found an association between internal microbiome diversity and social interactions in wild populations (Levin et al. 2016; Moeller et al. 2016; Tung et al. 2015). We found that dulling increased microbiome diversity by increasing the abundance of rare phyla, but the exact mechanism that resulted in these changes is still unclear. One possibility is that the altered social interactions associated with dulling might have resulted in different exposure to socially acquired microbes that subsequently influenced performance. For example, Levin et al. (2016) found that more close social interactions in barn swallows were associated with a more diverse internal microbiome. Alternatively, the differences in microbiome that we observed might have been a downstream consequence of changes in focal female behavior and physiology. Evidence in other species suggests that microbiome diversity might alter traits associated with performance, such as glucocorticoid regulation (Noguera et al. 2018), growth (Kohl et al. 2018), and immunity (Warne et al. 2019). Dulled females in our study also provisioned at a higher rate, and it is possible that exposure to more diverse food resources could have contributed to greater opportunities to acquire diverse environmental microbes. The design of our experiment does not allow us to distinguish between these possibilities at present and subsequent work that directly manipulates microbiome compositions under natural conditions will be illuminating.

Somewhat surprisingly, we did not find any evidence that plumage dulling—and subsequent changes in the social environment—influenced glucocorticoid regulation. Previous work showed that naturally bright females had a stronger corticosterone response to handling stress and were more resilient to natural stressors (Taff et al. 2019b). One potentially important difference is that the observational study was conducted in a year with extremely challenging weather conditions that resulted in a large percentage of nests failing (Taff et al. 2018). The extent to which glucocorticoid regulation predicts reproductive success in this population also varies with weather conditions (Vitousek et al. 2018a), so it is possible that under more challenging conditions signal manipulation and the experienced social environment might have interacted to influence performance. In this study, however, we found no evidence that changes in glucocorticoid regulation played an important role in coordination of the phenotypic changes that we observed. In contrast, we found that dulling resulted in an altered glucose response to handling. At present, it is unclear what caused this change, but if dulled females were changing their investment in foraging due to social interactions, that may have also resulted in changes to their energy reserves.

Moving forward, one of the challenges in understanding how dynamic feedback functions in signal evolution is to untangle the causal pathways that connect aspects of the phenotype at different timescales. Unlike honesty mechanisms that are essentially unidirectional, demonstrating causality in dynamic systems will likely require thorough time series data on multiple aspects of physiology and behavior (Taff & Vitousek 2016). Recent advances in bio-logging and remote sensing devices make it increasingly possible to collect extensive behavioral data under natural conditions (Krause et al. 2013), but comparable logging for most physiological measures is still lagging.

Regardless of the exact causal pathways that link signal expression to performance, our results demonstrate that bearing a particular signaling phenotype directly affects a suite of integrated traits. Thus, the apparent honesty of some signals might be an emergent property of feedback between physiology, signal expression, and the social environment rather than purely a result of intrinsic between-individual differences. In this case, individuals that are best able to flexibly coordinate various aspects of the phenotype in response to changing contexts and social conditions might achieve the highest fitness. It is important to note that this view is not incompatible with the specific unidirectional honesty mechanisms proposed by earlier work. Rather, we suggest that mechanisms suspected to play a role in signal production might act by setting the initial conditions on which feedback operates and be modified by social experience. Understanding how signal variation arises and is maintained will require integrative work that identifies specific mechanisms but also manipulates the signals and social environments in which those mechanisms operate.

## ACKNOWLEDGMENTS

We would like to thank David Winkler for establishing the research population used in this study and the many field and lab assistants who helped with data collection. CCT was funded by an Edward Rose postdoctoral fellowship from the Cornell Lab of Ornithology. Additional funding for this research was provided by DARPA D17AP00033 and NSF IOS 1457151 to MV. The views, opinions and/or findings expressed are those of the authors and should not be interpreted as representing the official views or policies of the Department of Defense or the U.S. Government.

