## Supplemental Figures and Tables for "Plumage manipulation alters the integration of social behavior, physiology, internal microbiome, and fitness"

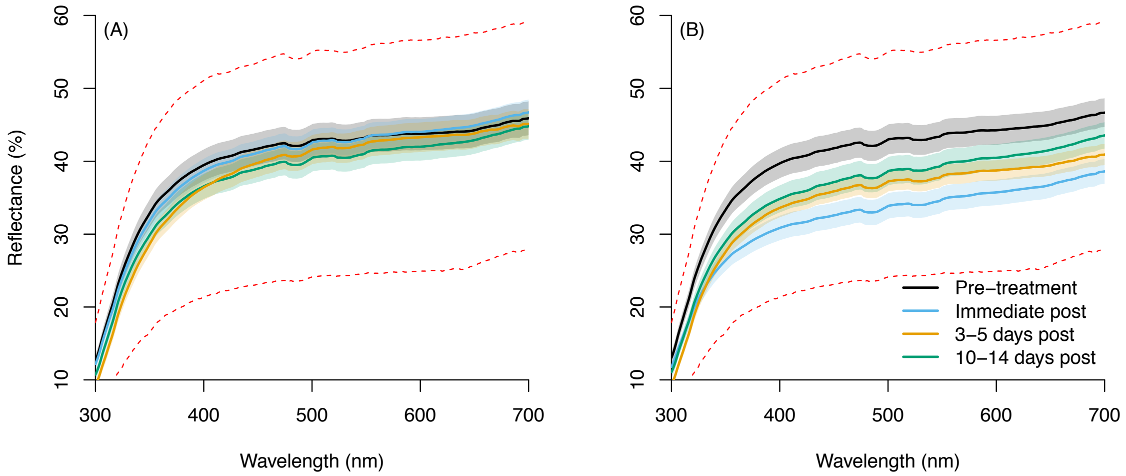


**Figure S1.** Effects of sham control (A) and dulling (B) treatments on breast plumage reflectance. Solid lines indicate the mean reflectance across all females for feathers collected pre-treatment (black), immediately post treatment (blue), 3-5 days post treatment (orange), or 10-14 days post treatment (green). Shaded regions illustrate the standard error of the mean for each measurement. Red dashed lines are the 5th and 95th percentiles for individual feather measurements.

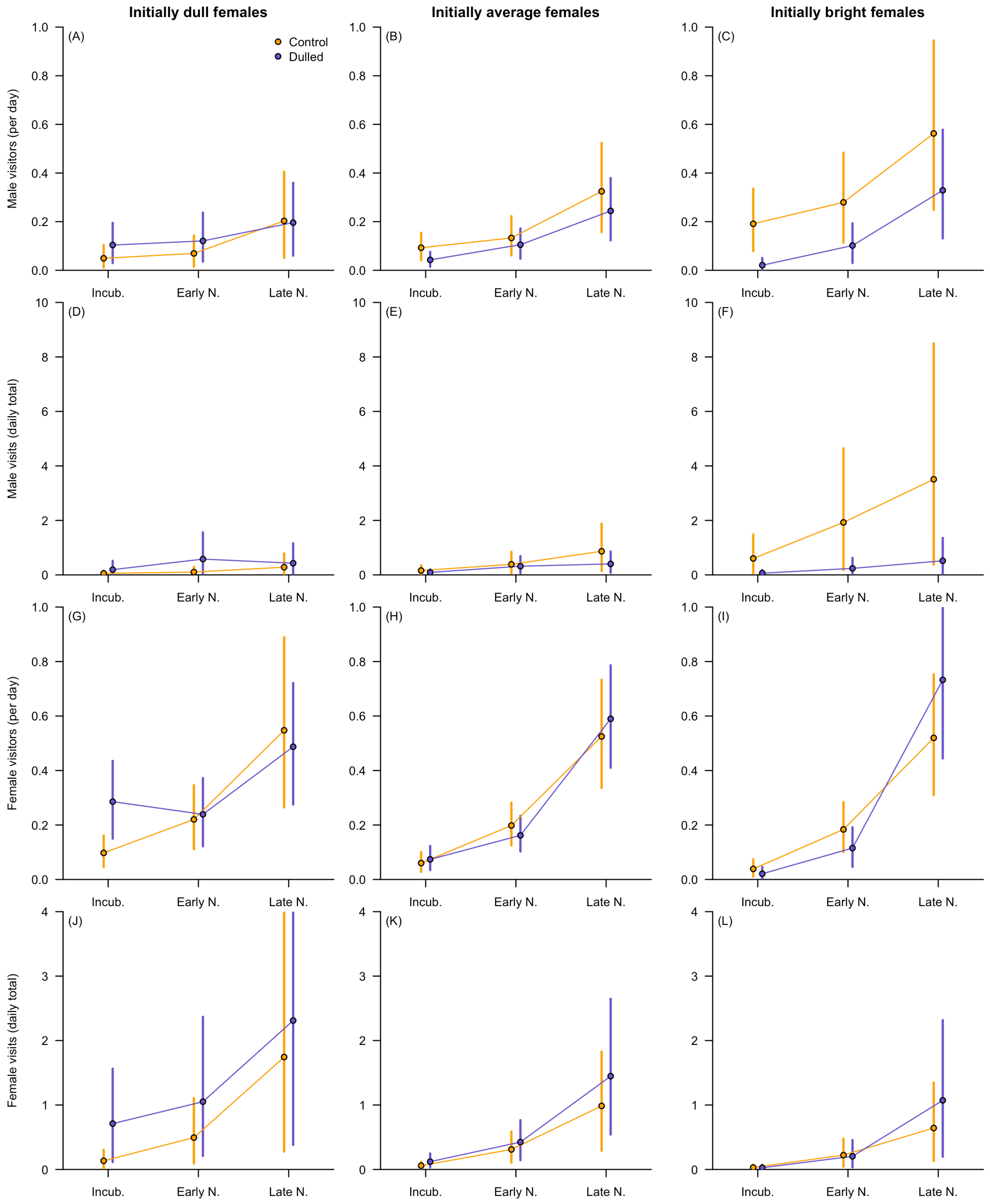


**Figure S2.** Model predicted relationship between treatment, breeding stage, initial plumage brightness and the number of total or unique visits by males and females based on the full models for each response variable in table S1. Columns illustrate the predicted relationship for females that initially had plumage 1 SD below the mean (left column), at the population mean (center column), or 1 SD above the mean (right column). Rows illustrate the predicted relationships for the number of unique male visitors (A-C), total male visitors (D-F), unique female visitors (G-I), or total female visitors (J-L). Within each panel, blue and orange points and lines indicate the predicted response value for dulled or control females with 95% confidence intervals during incubation, early nestling, and late nestling stages.

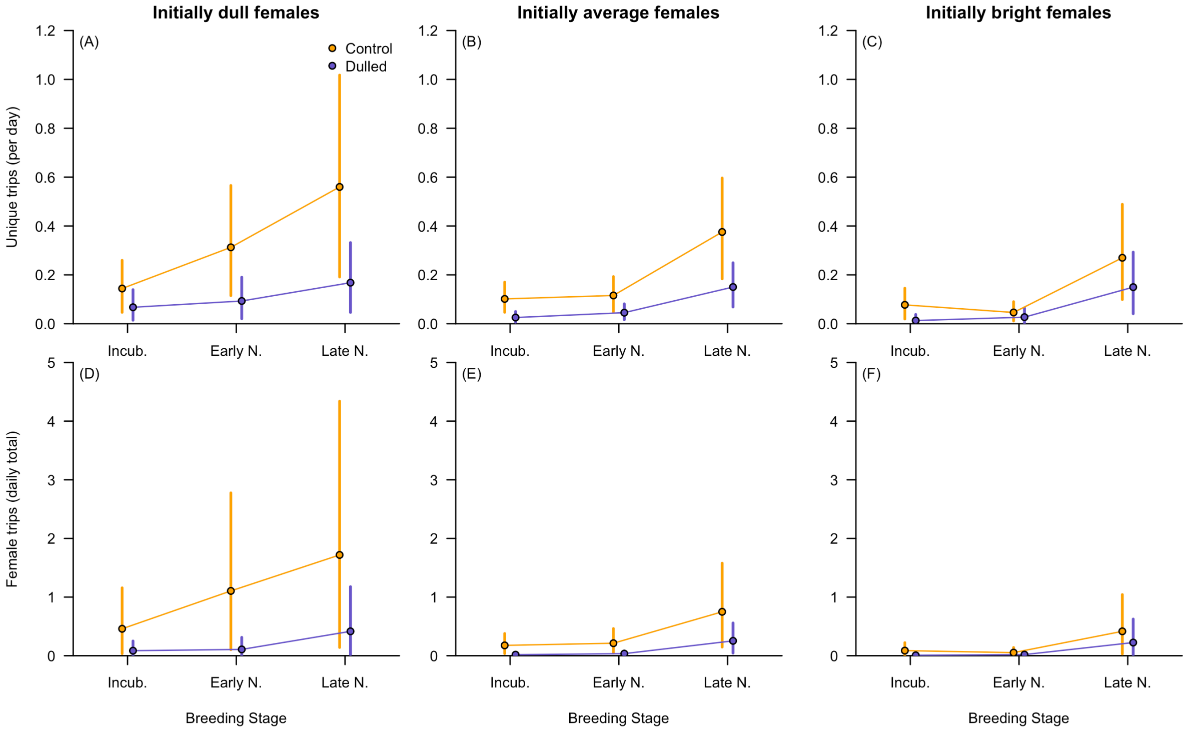


**Figure S3.** Model predicted relationship between treatment, breeding stage, initial plumage brightness and the number of total or unique trips made by focal females in this study to other monitored boxes. Columns are the predicted relationship for females that were initially 1 SD below the mean (left column), average (center column), or 1 SD above the mean (right column). Rows illustrate the predicted relationship for the number of trips made to unique boxes per day (A-C) and the total number of trips made per day (D-F).

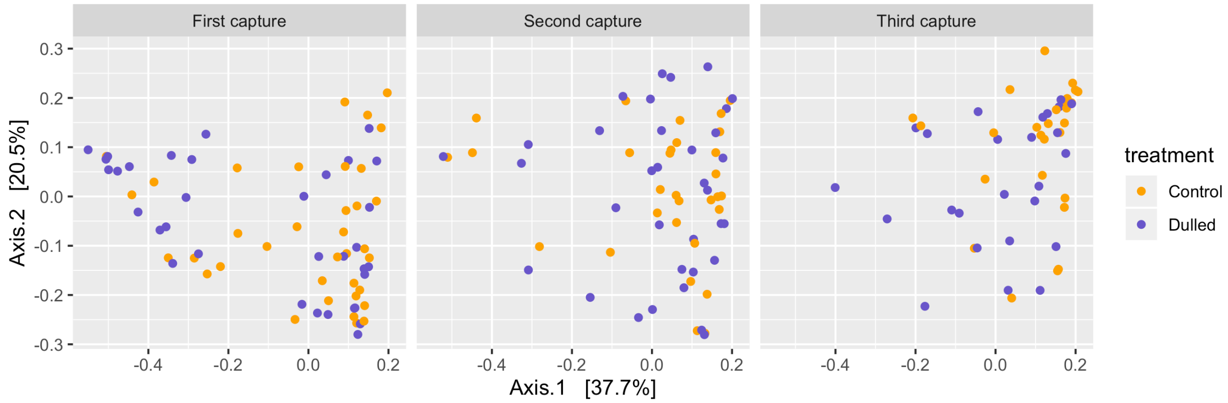


**Figure S4.** Nonmetric multidimensional scaling plots based on weighted unifrac distance comparing control (orange) and dulled (blue) birds at the first, second, and third capture.

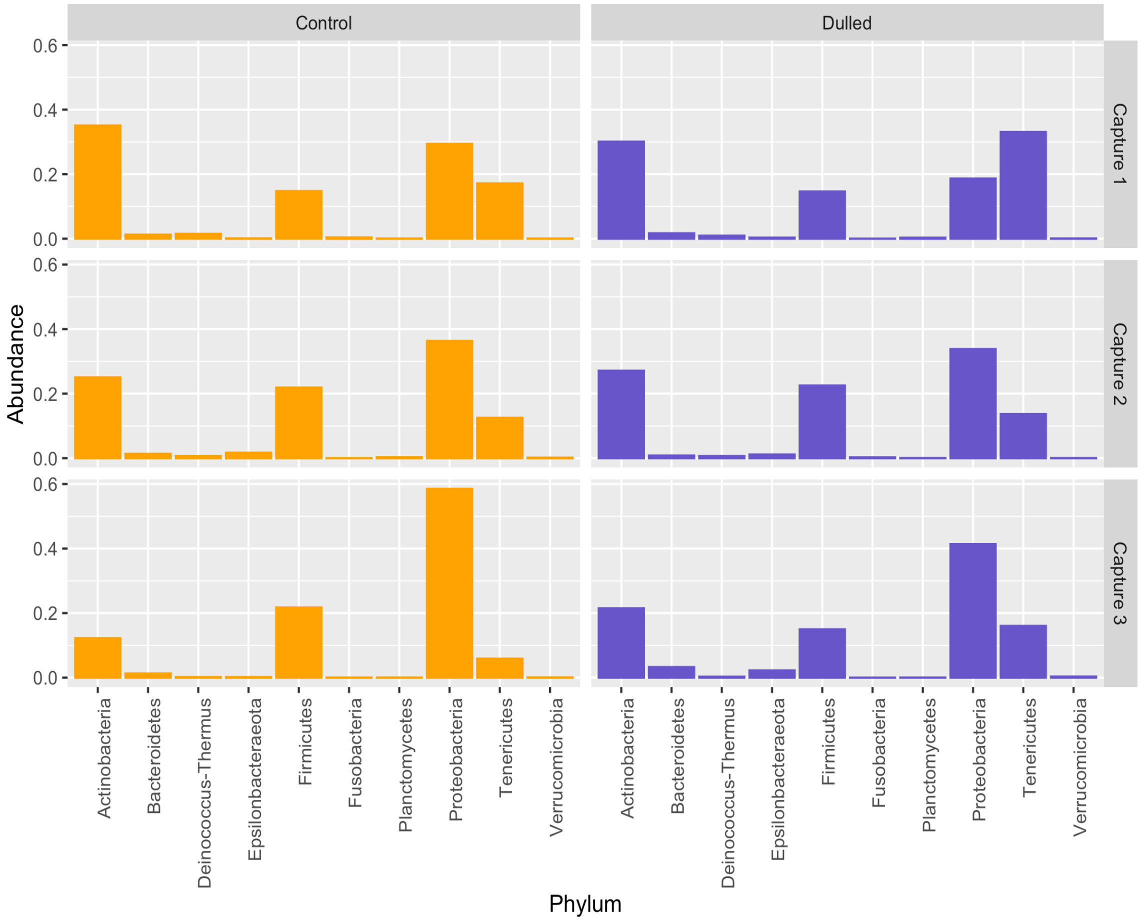


**Figure S5.** Relative abundance of sequences from different phyla by treatment group and capture number. Only phyla with average relative abundance > 0.1 % across all samples are shown.

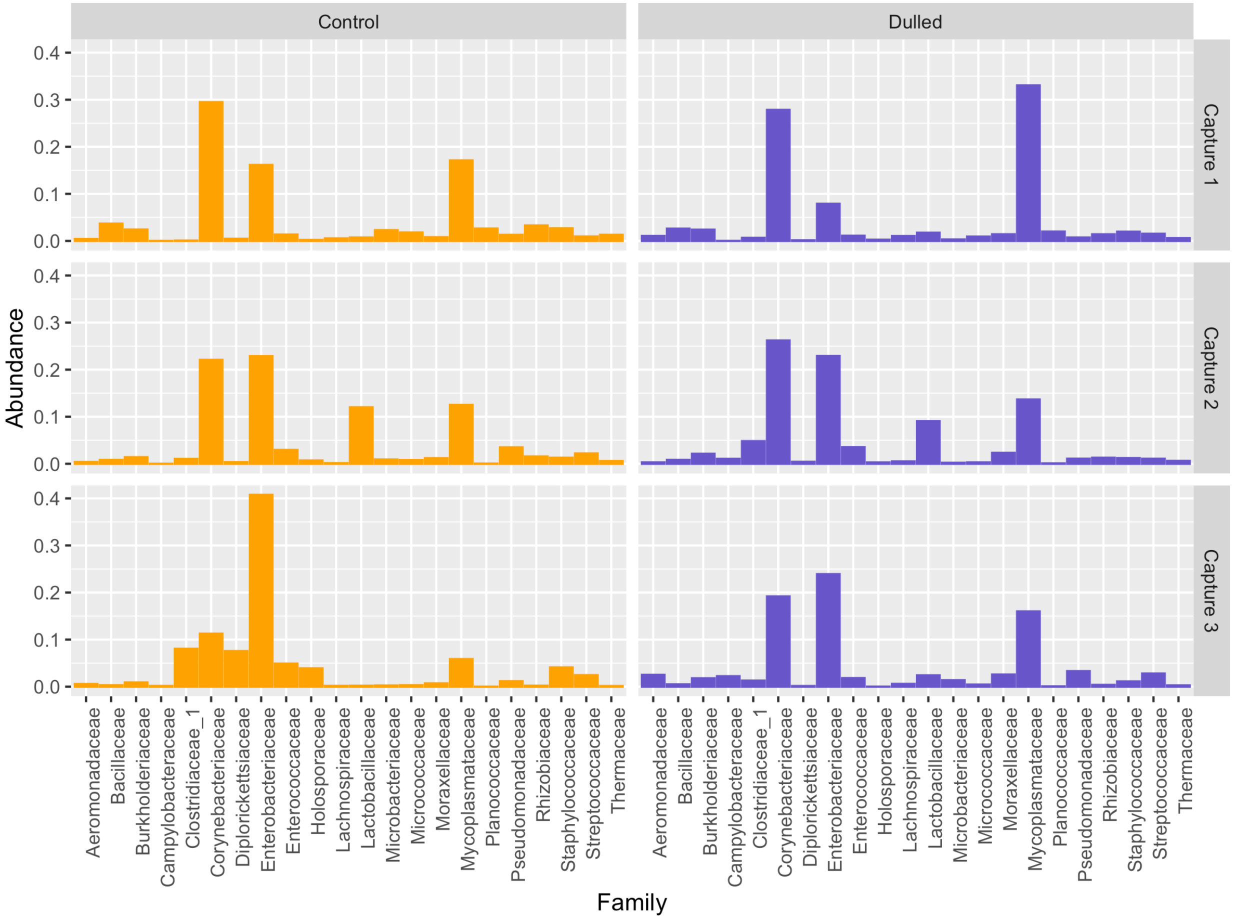


**Figure S6.** Relative abundance of different families across treatment groups and captures. Only families with average relative abundance values > 0.5% across all samples are plotted.

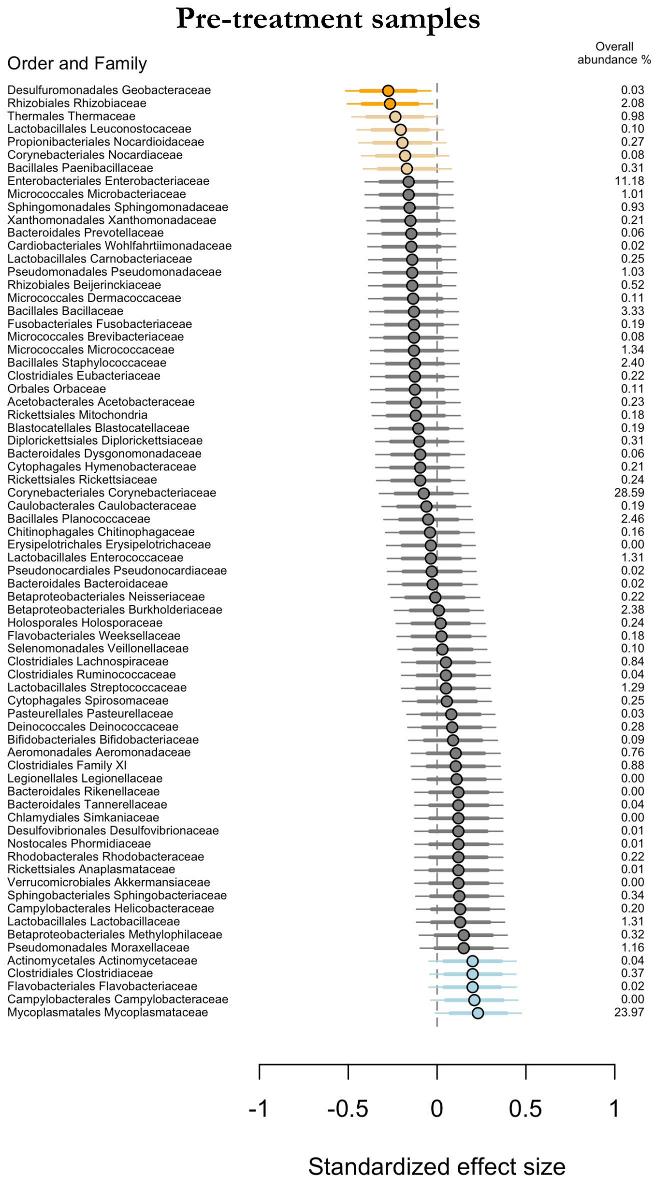


**Figure S7.** Family level comparisons of relative abundance for dulled and control samples collected prior to treatments. Points denote estimated effect sizes and lines denote the 95% (thin line) and 80% (thick line) confidence interval for effects. Colors indicate families where the 95% CI did not cross 0 (dark blue and dark orange), where the 80% CI did not cross 0 (light blue and light orange), and where both CIs crossed 0 (gray). Families in which dulled birds had higher relative abundance are illustrated in blue and those where control birds had higher relative abundance are in orange. Only families with average relative abundance across all samples > 0.05 % are shown.

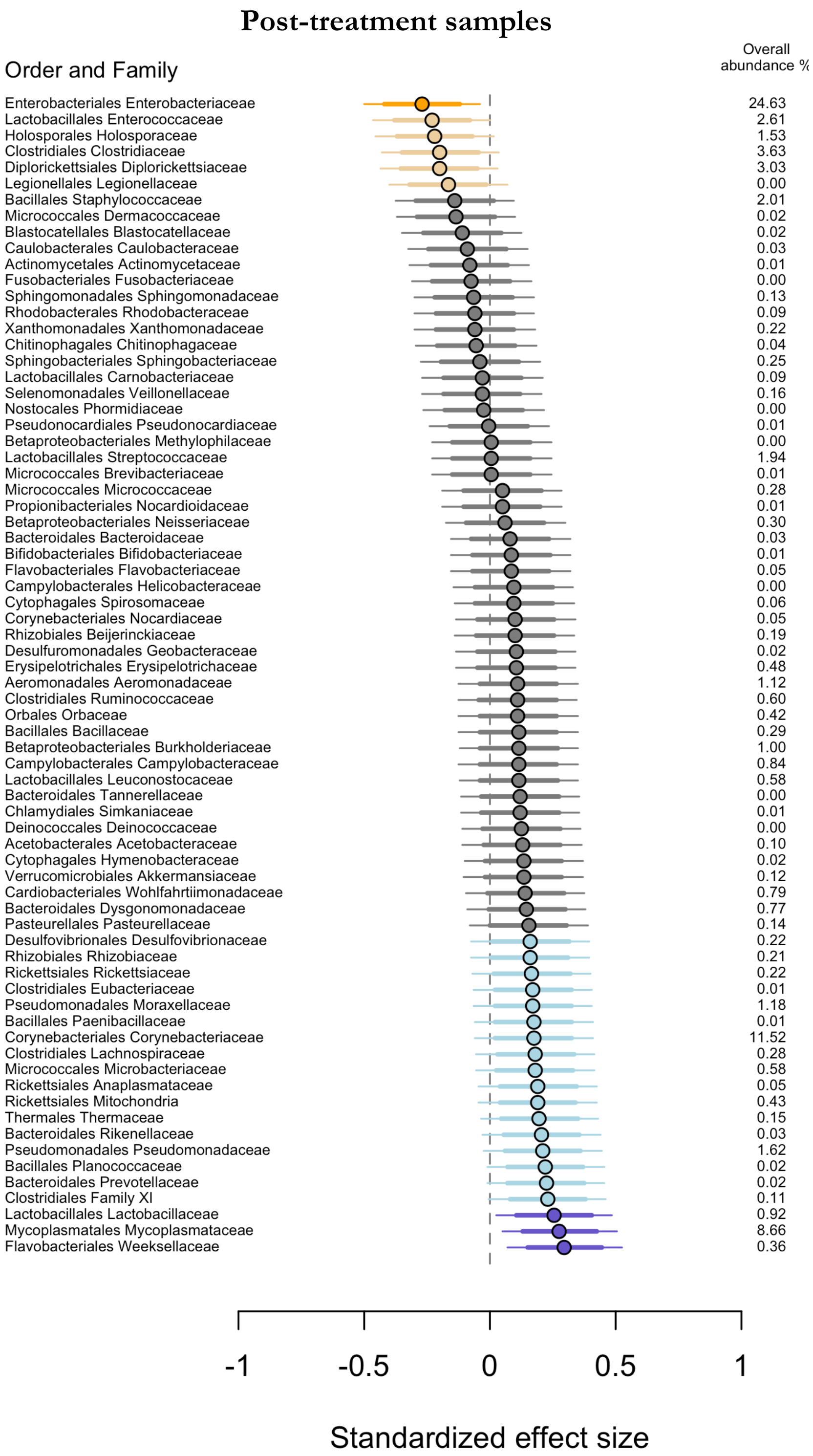


**Figure S8.** Family level comparisons of relative abundance for dulled and control samples collected after treatments. Points denote estimated effect sizes and lines denote the 95% (thin line) and 80% (thick line) confidence interval for effects. Colors indicate families where the 95% CI did not cross 0 (dark blue and dark orange), where the 80% CI did not cross 0 (light blue and light orange), and where both CIs crossed 0 (gray). Families in which dulled birds had higher relative abundance are illustrated in blue and those where control birds had higher relative abundance are in orange. Only families with average relative abundance across all samples > 0.05 % are shown.

**Table S1.** Model comparison for visits received by conspecific males and females at the focal box. All models include counts at the day level as the response variable; individual band and date in the season are included as random effects. All predictors listed in interactions were also included as main effects. Stage was entered as a three level factor for incubation, early nestling, and late nestling stages. Brightness is the initial breast brightness measure collected before treatments were applied. For zero-inflated negative binomials predictors were only included in the conditional part of the model.

| Response | Model | $\Delta AIC_c$ | Log Lik | *k* | Weight |
|---|---|---|---|---|---|
| *Total daily visits by females: zero-inflated negative binomial; n = 1,167 days; 63 focal boxes* | | | | | |
| | Treatment * Stage * Brightness | 0.0 | -1240.9 | 16 | 0.99 |
| | Treatment * Stage + Treatment * Brightness | 10.6 | -1250.3 | 12 | 0.01 |
| | Stage | 10.9 | -1255.6 | 7 | 0.00 |
| | Treatment * Brightness + Stage | 11.4 | -1252.7 | 10 | 0.00 |
| | Intercept Only | 80.5 | -1292.4 | 5 | 0.00 |
| *Unique daily female visitors: Poisson; n = 1,167 days; 63 focal boxes* | | | | | |
| | Treatment * Stage * Brightness | 0.0 | -734.1 | 14 | 1.00 |
| | Stage | 31.6 | -759.0 | 5 | 0.00 |
| | Treatment * Stage + Treatment * Brightness | 33.9 | -755.1 | 10 | 0.00 |
| | Treatment * Brightness + Stage | 36.0 | -758.2 | 8 | 0.00 |
| | Intercept Only | 96.4 | -793.4 | 3 | 0.00 |
| *Total daily visits by males: zero-inflated negative binomial; n = 1,167 days; 63 focal boxes* | | | | | |
| | Treatment * Stage * Brightness | 0.0 | -1336.4 | 16 | 0.70 |
| | Treatment * Stage + Treatment * Brightness | 2.9 | -1341.9 | 12 | 0.17 |
| | Treatment * Brightness + Stage | 4.0 | -1344.5 | 10 | 0.10 |
| | Stage | 6.1 | -1348.6 | 7 | 0.03 |
| | Intercept Only | 80.2 | -1387.7 | 5 | 0.00 |
| *Unique daily male visitors: Poisson; n = 1,167 days; 63 focal boxes* | | | | | |
| | Treatment * Brightness + Stage | 0.0 | -627.1 | 8 | 0.38 |
| | Treatment * Stage * Brightness | 0.1 | -621.0 | 14 | 0.37 |
| | Stage | 1.5 | -630.9 | 5 | 0.18 |
| | Treatment * Stage + Treatment * Brightness | 3.4 | -626.8 | 10 | 0.07 |
| | Intercept Only | 40.4 | -652.4 | 3 | 0.00 |

**Table S2.** Details of best-supported models for conspecific visitation rates.

| Response | Model | Estimate | 2.5 % CI | *97.5 % CI* |
|---|---|---|---|---|
| *Total daily visits by females: zero-inflated negative binomial; n = 1,167 days; 63 focal boxes* | | | | |
| | Intercept (Control; Incubation) | **-2.96** | **-3.94** | **-1.98** |
| | Dulled | 0.74 | -0.60 | 2.08 |
| | Initial Brightness | **-0.76** | **-1.52** | **-0.01** |
| | Early Nestlings | **1.70** | **1.00** | **2.41** |
| | Late Nestlings | **2.89** | **2.12** | **3.59** |
| | Dulled * Initial Brightness | -0.96 | -2.20 | 0.27 |
| | Dulled * Early Nestlings | -0.41 | -1.40 | 0.58 |
| | Dulled * Late Nestlings | -0.33 | -1.33 | 0.67 |
| | Initial Brightness * Early Nestlings | 0.38 | -0.06 | 0.81 |
| | Initial Brightness * Late Nestlings | 0.29 | -0.28 | 0.86 |
| | Dulled * Brightness * Early Nestlings | 0.53 | -0.25 | 1.30 |
| | Dulled * Brightness * Late Nestlings | **1.06** | **0.14** | **1.97** |
| *Unique daily female visitors: Poisson; n = 1,167 days; 63 focal boxes* | | | | |
| | Intercept (Control; Incubation) | **-2.86** | **-3.49** | **-2.23** |
| | Dulled | 0.20 | -0.66 | 1.07 |
| | Initial Brightness | **-0.48** | **-0.90** | **-0.07** |
| | Early Nestlings | **1.22** | **0.58** | **1.86** |
| | Late Nestlings | **2.20** | **1.56** | **2.84** |
| | Dulled * Initial Brightness | -0.88 | -1.63 | -0.14 |
| | Dulled * Early Nestlings | -0.41 | -1.30 | 0.49 |
| | Dulled * Late Nestlings | -0.08 | -0.93 | 0.77 |
| | Initial Brightness * Early Nestlings | 0.40 | -0.05 | 0.84 |
| | Initial Brightness * Late Nestlings | 0.47 | -0.01 | 0.95 |
| | Dulled * Brightness * Early Nestlings | 0.60 | -0.18 | 1.39 |
| | Dulled * Brightness * Late Nestlings | **1.11** | **0.34** | **1.88** |
| *Total daily visits by males: zero-inflated negative binomial; n = 1,167 days; 63 focal boxes* | | | | |
| | Intercept (Control; Incubation) | **-1.99** | **-3.08** | **-0.89** |
| | Dulled | -0.55 | -2.07 | 0.97 |
| | Initial Brightness | **1.25** | **0.20** | **2.29** |
| | Early Nestlings | **0.89** | **0.53** | **1.24** |
| | Late Nestlings | **1.69** | **1.23** | **2.08** |
| | Dulled * Initial Brightness | **-1.82** | **-3.39** | **-0.25** |
| | Dulled * Early Nestlings | 0.37 | -0.29 | 1.03 |
| | Dulled * Late Nestlings | -0.19 | -0.80 | 0.42 |
| | Initial Brightness * Early Nestlings | 0.28 | -0.13 | 0.69 |
| | Initial Brightness * Late Nestlings | 0.07 | -0.32 | 0.46 |
| | Dulled * Brightness * Early Nestlings | -0.13 | -0.81 | 0.56 |
| | Dulled * Brightness * Late Nestlings | 0.62 | -0.08 | 1.33 |
| *Unique daily male visitors: Poisson; n = 1,167 days; 63 focal boxes* | | | | |
| | Intercept (Control: Incubation) | -2.51 | -3.12 | -1.91 |
| | Dulled | -0.24 | -0.94 | 0.47 |
| | Initial Brightness | **0.64** | **0.13** | **1.16** |
| | Early Nestlings | **0.51** | **0.09** | **0.93** |
| | Late Nestlings | **1.31** | **0.94** | **1.69** |
| | Dulled * Initial Brightness | -0.63 | -1.35 | 0.09 |
| *Unique daily male visitors: Poisson; n = 1,167 days; 63 focal boxes (delta $AIC_c$ = 0.1)* | | | | |
| | Intercept (Control; Incubation) | -2.43 | -3.06 | -1.80 |
| | Dulled | -0.80 | -1.75 | 0.16 |
| | Initial Brightness | **0.71** | **0.11** | **1.30** |
| | Early Nestlings | **0.36** | **-0.20** | **0.93** |
| | Late Nestlings | **1.26** | **0.75** | **1.77** |
| | Dulled * Initial Brightness | **-1.57** | **-2.58** | **-0.57** |
| | Dulled * Early Nestlings | 0.56 | -0.30 | 1.43 |
| | Dulled * Late Nestlings | 0.52 | -0.29 | 1.33 |
| | Initial Brightness * Early Nestlings | 0.02 | -0.52 | 0.55 |
| | Initial Brightness * Late Nestlings | -0.17 | -0.67 | 0.32 |
| | Dulled * Brightness * Early Nestlings | 0.77 | -0.18 | 1.71 |
| | Dulled * Brightness * Late Nestlings | **1.31** | **0.40** | **2.21** |

**Table S3.** Model comparison table for social mate activity and provisioning rate at the focal box. Response variables are the standardized number of total RFID reads or standardized number of total provisioning trips made each day. All models include nest box identity and date in the season as random effects.

| Response Model | $\Delta AIC_c$ | Log Lik | *k* | Weight |
|---|---|---|---|---|
| *Total daily nest box attendance by males: LMM, n = 487 days for 39 nest boxes* | | | | |
| Nestling Age + Nestling Age$^2$ | 0.0 | -598.1 | 6 | 0.77 |
| Treatment + Nestling Age + Nestling Age$^2$ | 2.7 | -598.4 | 7 | 0.21 |
| Intercept Only | 8.3 | -604.3 | 4 | 0.01 |
| Treatment * Brightness + Nestling Age + Nestling Age$^2$ | 8.8 | -599.37 | 9 | 0.01 |
| *Total daily provisioning trips by males: LMM, n = 456 days for 37 males* | | | | |
| Nestling Age + Nestling Age$^2$ | 0.0 | -445.6 | 6 | 0.85 |
| Treatment + Nestling Age + Nestling Age$^2$ | 3.5 | -446.3 | 7 | 0.15 |
| Treatment * Brightness + Nestling Age + Nestling Age$^2$ | 11.2 | -448.1 | 9 | 0.00 |
| Intercept Only | 169.8 | -532.5 | 4 | 0.00 |

**Table S4.** Model comparison for focal female activity and provisioning rate at her own box. Response variables are the standardized number of total RFID reads or standardized number of total provisioning trips made each day. Each model includes box identity and date in the season as a random effect.

| Response Model | $\Delta AIC_c$ | Log Lik | *k* | Weight |
|---|---|---|---|---|
| *Total daily nest box attendance by focal females: LMM, n = 752 days for 59 focal females* | | | | |
| Intercept Only | 0.0 | -966.9 | 4 | 1.00 |
| Nestling Age + Nestling Age$^2$ | 13.1 | -971.4 | 6 | 0.00 |
| Treatment + Nestling Age + Nestling Age$^2$ | 16.3 | -972.0 | 7 | 0.00 |
| Treatment * Brightness + Nestling Age + Nestling Age$^2$ | 25.1 | -974.3 | 9 | 0.00 |
| *Total daily provisioning trips by focal females: LMM, n = 752 days for 59 focal females* | | | | |
| Treatment + Nestling Age + Nestling Age$^2$ | 0.0 | -851.6 | 7 | 0.76 |
| Nestling Age + Nestling Age$^2$ | 2.7 | -854.0 | 6 | 0.19 |
| Treatment * Brightness + Nestling Age + Nestling Age$^2$ | 5.6 | -852.4 | 9 | 0.05 |
| Intercept Only | 36.8 | -873.1 | 4 | 0.00 |

**Table S5.** Summary of best-supported model for female provisioning rate. The response variable is the standardized number of total daily provisioning trips. Female identity and date are included as random effects.

| Response Model | Estimate | 2.5 % CI | *97.5 % CI* |
|---|---|---|---|
| *Total daily provisioning trips by focal females* | | | |
| Intercept (Control) | **-1.24** | **-1.58** | **-0.90** |
| Dulled | **0.35** | **0.09** | **0.60** |
| Nestling Age | **.12** | **0.07** | **0.17** |
| Nestling Age$^2$ | -0.0 | -0.0 | 0.0 |

**Table S6.** Model selection table for daily focal female trips to other monitored boxes.

| Response Model | $\Delta AIC_c$ | Log Lik | *k* | Weight |
|---|---|---|---|---|
| *Total daily trips by focal females: zero-inflated negative binomial; n = 1,223 days; 64 focal females* | | | | |
| Treatment * Stage * Brightness | 0.0 | -931.1 | 16 | 0.99 |
| Treatment * Stage + Treatment * Brightness | 10.6 | -940.5 | 12 | 0.01 |
| Treatment * Brightness + Stage | 13.1 | -943.8 | 10 | 0.00 |
| Stage | 15.9 | -948.2 | 7 | 0.00 |
| Intercept Only | 72.1 | -978.3 | 5 | 0.00 |
| *Unique daily trips to other boxes by focal females: Poisson GLMM, n = 1,223 days for 64 focal females* | | | | |
| Treatment * Stage * Brightness | 0.0 | -580.9 | 14 | 1.00 |
| Treatment * Brightness + Stage | 13.4 | -593.7 | 8 | 0.00 |
| Treatment * Stage + Treatment * Brightness | 16.6 | -593.3 | 10 | 0.00 |
| Stage | 18.7 | -599.4 | 5 | 0.00 |
| Intercept Only | 54.4 | -619.2 | 3 | 0.00 |

**Table S7.** Details for best-supported models for total trips and unique trips to other boxes made by focal females.

| Response Model | Estimate | 2.5 % CI | *97.5 % CI* |
|---|---|---|---|
| *Total daily trips by focal females: zero-inflated negative binomial; n = 1,223 days; 64 focal females* | | | |
| Intercept (Control; Incubation) | **-1.88** | **-2.94** | **-0.83** |
| Dulled | **-2.39** | **-4.08** | **-0.70** |
| Initial Brightness | -0.83 | -1.82 | 0.16 |
| Early Nestlings | 0.19 | -0.29 | 0.67 |
| Late Nestlings | **1.45** | **0.87** | **2.03** |
| Dulled * Initial Brightness | -0.58 | -2.45 | 1.29 |
| Dulled * Early Nestlings | 0.56 | -0.46 | 1.58 |
| Dulled * Late Nestlings | **1.30** | **0.23** | **2.36** |
| Initial Brightness * Early Nestlings | **-0.70** | **-1.28** | **-0.13** |
| Initial Brightness * Late Nestlings | 0.14 | -0.47 | 0.74 |
| Dulled * Brightness * Early Nestlings | 1.21 | -0.22 | 2.64 |
| Dulled * Brightness * Late Nestlings | 0.98 | -0.39 | 2.34 |
| *Unique daily trips to other boxes by focal females: Poisson GLMM, n = 1,223 days for 64 focal females* | | | |
| Intercept (Control; Incubation) | **-2.34** | **-2.97** | **-1.70** |
| Dulled | **-1.45** | **-2.53** | **-0.37** |
| Initial Brightness | -0.32 | -0.86 | 0.22 |
| Early Nestlings | 0.13 | -0.39 | 0.65 |
| Late Nestlings | **1.32** | **0.79** | **1.85** |
| Dulled * Initial Brightness | -0.63 | -1.84 | 0.58 |
| Dulled * Early Nestlings | 0.49 | -0.48 | 1.46 |
| Dulled * Late Nestlings | 0.53 | -0.39 | 1.45 |
| Initial Brightness * Early Nestlings | **-0.66** | **-1.02** | **-0.29** |
| Initial Brightness * Late Nestlings | -0.04 | -0.43 | 0.35 |
| Dulled * Brightness * Early Nestlings | 0.94 | -0.25 | 2.13 |
| Dulled * Brightness * Late Nestlings | 0.94 | -0.14 | 2.01 |

**Table S8.** Model selection tables for focal female physiological measures. For each response, we fit the same set of four linear models. Sample sizes vary between response variables because not all measures were taken for all females or because nests failed before samples could be collected. In all models 'initial' refers to the pre-treatment measure of the response variable.

| Response Model | $\Delta AIC_c$ | Log Lik | *k* | Weight |
|---|---|---|---|---|
| *Baseline corticosterone at second capture; n = 64* | | | | |
| Intercept Only | 0.0 | -87.9 | 2 | 0.66 |
| Initial | 2.2 | -87.9 | 3 | 0.22 |
| Treatment + Initial | 3.8 | -87.6 | 4 | 0.10 |
| Treatment * Brightness + Initial | 7.8 | -87.6 | 6 | 0.01 |
| *Stress-induced corticosterone at second capture; n = 64* | | | | |
| Initial | 0.0 | -88.7 | 3 | 0.53 |
| Intercept Only | 2.0 | -90.8 | 2 | 0.20 |
| Treatment + Initial | 2.3 | -88.7 | 4 | 0.17 |
| Treatment * Brightness + Initial | 3.4 | -86.9 | 6 | 0.10 |
| *Dexamethasone-induced corticosterone at second capture; n = 62* | | | | |
| Initial | 0.0 | -85.7 | 3 | 0.44 |
| Treatment + Initial | 0.9 | -85.0 | 4 | 0.28 |
| Treatment * Brightness + Initial | 1.7 | -83.0 | 6 | 0.19 |
| Intercept Only | 3.3 | -88.4 | 2 | 0.09 |
| *Baseline glucose at second capture; n = 62* | | | | |
| Initial | 0.0 | -85.7 | 3 | 0.60 |
| Treatment + Initial | 2.1 | -85.6 | 4 | 0.21 |
| Intercept Only | 3.1 | -88.3 | 2 | 0.13 |
| Treatment * Brightness + Initial | 4.9 | -84.5 | 6 | 0.05 |
| *Stress-induced glucose at second capture; n = 65* | | | | |
| Treatment * Brightness + Initial | 0.0 | -83.3 | 6 | 0.70 |
| Treatment + Initial | 3.0 | -87.2 | 4 | 0.16 |
| Initial | 4.2 | -88.9 | 3 | 0.09 |
| Intercept Only | 4.9 | -90.4 | 2 | 0.06 |
| *Mass at second capture; n = 65* | | | | |
| Initial | 0.0 | -84.7 | 3 | 0.72 |
| Treatment + Initial | 2.2 | -84.6 | 4 | 0.25 |
| Treatment * Brightness + Initial | 6.6 | -84.4 | 6 | 0.03 |
| Intercept Only | 12.6 | -92.1 | 2 | 0.00 |
| *Baseline corticosterone at third capture; n = 50* | | | | |
| Intercept Only | 0.0 | -70.9 | 2 | 0.34 |
| Initial | 0.1 | -69.9 | 3 | 0.32 |
| Treatment + Initial | 0.3 | -68.8 | 4 | 0.30 |
| Treatment * Brightness + Initial | 3.9 | -68.0 | 6 | 0.05 |
| *Mass at third capture; n = 51* | | | | |
| Intercept Only | 0.0 | -72.3 | 2 | 0.51 |
| Initial | 0.7 | -71.5 | 3 | 0.36 |
| Treatment + Initial | 2.9 | -71.5 | 4 | 0.12 |
| Treatment * Brightness + Initial | 7.8 | -71.4 | 6 | 0.01 |

**Table S9.** Details for supported models of focal female physiology that included a treatment effect.

| Response Model | Estimate | 2.5 % CI | *97.5 % CI* |
|---|---|---|---|
| *Stress-induced glucose at second capture; n = 65* | | | |
| Intercept (Control) | **-0.22** | **-0.11** | **0.55** |
| Treatment | **-0.50** | **-0.98** | **-0.03** |
| Brightness | 0.03 | -0.27 | 0.32 |
| Treatment * Brightness | **0.47** | **0.01** | **0.93** |

**Table S10.** Model selection table for measures of microbiome diversity with reads rarefied to 153. Model rankings with a rarefaction limit of 1500 were similar, except that for Faith's D at third capture the intercept only model had the lowest $AIC_c$ value.

| Response Model | $\Delta AIC_c$ | Log Lik | *k* | Weight |
|---|---|---|---|---|
| *Simpson diversity at second capture; n = 63* | | | | |
| Intercept Only | 0.0 | -68.9 | 4 | 0.78 |
| Initial | 2.0 | -89.2 | 3 | 0.21 |
| Treatment + Initial | 2.2 | -88.2 | 4 | 0.19 |
| Treatment * Brightness + Initial | 6.4 | -87.9 | 6 | 0.02 |
| *Simpson diversity at third capture; n = 52* | | | | |
| Treatment + Initial | 0.0 | -68.9 | 4 | 0.78 |
| Treatment * Brightness + Initial | 4.4 | -68.5 | 6 | 0.09 |
| Initial | 4.6 | -72.3 | 3 | 0.08 |
| Initial | 5.1 | -73.7 | 2 | 0.06 |
| *Shannon diversity at second capture; n = 63* | | | | |
| Intercept Only | 0.0 | -89.3 | 2 | 0.51 |
| Treatment + Initial | 1.3 | -87.8 | 4 | 0.26 |
| Initial | 2.0 | -89.3 | 3 | 0.18 |
| Treatment * Brightness + Initial | 4.6 | -87.0 | 6 | 0.05 |
| *Shannon diversity at third capture; n = 52* | | | | |
| Treatment + Initial | 0.0 | -69.5 | 4 | 0.71 |
| Intercept Only | 3.6 | -73.6 | 2 | 0.12 |
| Treatment * Brightness + Initial | 3.8 | -68.9 | 6 | 0.10 |
| Initial | 4.5 | -72.9 | 3 | 0.07 |
| *Faith's D at second capture; n = 63* | | | | |
| Treatment + Initial | 0.0 | -87.1 | 4 | 0.35 |
| Intercept Only | 0.1 | -89.4 | 2 | 0.33 |
| Initial | 1.1 | -88.8 | 3 | 0.20 |
| Treatment * Brightness + Initial | 2.1 | -85.7 | 6 | 0.12 |
| *Faith's D at third capture; n = 52* | | | | |
| Treatment + Initial | 0.0 | -68.5 | 4 | 0.77 |
| Treatment * Brightness + Initial | 3.1 | -67.6 | 6 | 0.16 |
| Intercept Only | 5.6 | -73.6 | 2 | 0.05 |
| Initial | 7.8 | -73.6 | 3 | 0.02 |

**Table S11.** Details for best-supported models of microbiome diversity at the third capture.

| Response Model | Estimate | 2.5 % CI | *97.5 % CI* |
|---|---|---|---|
| *Simpson diversity at third capture; n = 52* | | | |
| Intercept (Control) | -0.32 | -0.67 | 0.04 |
| Treatment | **0.70** | **0.17** | **1.22** |
| Initial Brightness | -0.20 | -0.45 | 0.06 |
| *Shannon diversity at third capture; n = 52* | | | |
| Intercept (Control) | -0.31 | -0.67 | 0.05 |
| Treatment | **0.70** | **0.16** | **1.24** |
| Initial Brightness | -0.11 | -0.38 | 0.15 |
| *Faith's D at third capture; n = 52* | | | |
| Intercept (Control) | -0.36 | -0.73 | -0.02 |
| Treatment | **0.84** | **0.32** | **1.36** |
| Initial Brightness | 0.04 | -0.24 | 0.33 |

**Table S12.** Model selection tables for relative abundance of particular phyla.

| Response Model | $\Delta AIC_c$ | Log Lik | *k* | Weight |
|---|---|---|---|---|
| *Relative abundance of Actinobacterial at third capture; n = 52* | | | | |
| Treatment | 0.0 | 19.0 | 3 | 0.53 |
| Intercept Only | 1.9 | 17.0 | 2 | 0.21 |
| Treatment + Initial | 2.2 | 19.1 | 4 | 0.18 |
| Initial | 4.0 | 17.0 | 3 | 0.07 |
| *Relative abundance of Bacteroidetes at third capture; n = 52* | | | | |
| Treatment | 0.0 | 95.8 | 3 | 0.53 |
| Intercept Only | 1.8 | 93.8 | 2 | 0.22 |
| Treatment + Initial | 2.2 | 95.9 | 4 | 0.18 |
| Initial | 3.8 | 93.9 | 3 | 0.08 |
| *Relative abundance of Firmicutes at third capture; n = 52* | | | | |
| Initial | 0.0 | 19.1 | 3 | 0.61 |
| Treatment + Initial | 1.7 | 19.7 | 4 | 0.36 |
| Intercept Only | 6.8 | 14.6 | 2 | 0.02 |
| Treatment | 7.8 | 15.2 | 3 | 0.01 |
| *Relative abundance of Proteobacteria at third capture; n = 52* | | | | |
| Treatment | 0.0 | 0.5 | 3 | 0.56 |
| Treatment + Initial | 1.7 | 0.9 | 4 | 0.24 |
| Intercept Only | 3.1 | -2.1 | 2 | 0.12 |
| Initial | 4.0 | -1.5 | 3 | 0.08 |
| *Relative abundance of Tenericutes at third capture; n = 52* | | | | |
| Treatment | 0.0 | 34.5 | 3 | 0.57 |
| Intercept Only | 2.3 | 32.2 | 2 | 0.18 |
| Treatment + Initial | 2.3 | 34.5 | 4 | 0.18 |
| Initial | 4.3 | 32.3 | 3 | 0.07 |

**Table S13.** Details of best-supported models for relative abundance of different phyla.

| Response Model | Estimate | 2.5 % CI | *97.5 % CI* |
|---|---|---|---|
| *Relative abundance of Actinobacteria at third capture; n = 52* | | | |
| Intercept (Control) | 0.12 | 0.06 | 0.19 |
| Treatment | **0.10** | **> 0.00** | **0.19** |
| *Relative abundance of Bacteroidetes at third capture; n = 52* | | | |
| Intercept (Control) | 0.01 | 0.01 | 0.02 |
| Treatment | **0.02** | **> 0.00** | **0.04** |
| *Relative abundance of Proteobacteria at third capture; n = 52* | | | |
| Intercept (Control) | 0.59 | 0.49 | 0.68 |
| Treatment | **-0.16** | **-0.29** | **-0.02** |
| *Relative abundance of Tenericutes at third capture; n = 52* | | | |
| Intercept (Control) | 0.06 | 0.01 | 0.11 |
| Treatment | **0.08** | **0.01** | **0.15** |

**Table S14.** Candidate model sets for nestling survival and phenotype. All models are binomial GLMMs with nest identity included as a random effect.

| Response | Model | $\Delta AIC_c$ | Log Lik | *k* | Weight |
|---|---|---|---|---|---|
| *Hatching success of nestlings; binomial GLMM; n = 365 individuals from 69 nests* | | | | | |
| | Treatment | 0.0 | -129.6 | 3 | 0.47 |
| | Treatment * Brightness | 0.2 | -127.6 | 5 | 0.42 |
| | Intercept Only | 2.9 | -132.0 | 2 | 0.11 |
| *Survival to day 12 for nestlings; binomial GLMM; n = 365 individuals from 69 nests* | | | | | |
| | Treatment * Brightness | 0.0 | -184.2 | 5 | 0.63 |
| | Treatment | 1.3 | -186.8 | 3 | 0.33 |
| | Intercept Only | 5.5 | -190.0 | 2 | 0.04 |
| *Survival to fledge for nestlings; binomial GLMM; n = 365 individuals from 69 nests* | | | | | |
| | Treatment * Brightness | 0.0 | -191.1 | 5 | 0.46 |
| | Treatment | 0.6 | -193.5 | 3 | 0.35 |
| | Intercept Only | 1.8 | -195.1 | 2 | 0.19 |
| *Nestling mass on day 12; LMM; n = 220 individuals from 53 nests* | | | | | |
| | Intercept Only | 0.0 | -244.7 | 3 | 0.78 |
| | Treatment | 2.6 | -245.0 | 4 | 0.21 |
| | Treatment * Brightness | 8.7 | -245.9 | 6 | 0.01 |
| *Nestling wing length on day 12; LMM; n = 222 individuals from 53 nests* | | | | | |
| | Intercept Only | 0.0 | -227.3 | 3 | 0.78 |
| | Treatment | 2.7 | -227.6 | 4 | 0.20 |
| | Treatment * Brightness | 8.1 | -228.2 | 6 | 0.01 |
| *Nestling bill + head on day 12; LMM; n = 221 individuals from 53 nests* | | | | | |
| | Intercept Only | 0.0 | -267.2 | 3 | 0.75 |
| | Treatment | 2.3 | -267.2 | 4 | 0.24 |
| | Treatment * Brightness | 8.4 | -268.2 | 6 | 0.01 |

**Table S15.** Details of best-supported models for nestling survival. Models with $\Delta AIC_c < 2$ are included.

| Response Model | Estimate | 2.5 % CI | *97.5 % CI* |
|---|---|---|---|
| *Hatching success of nestlings; GLMM; n = 365 individuals from 69 nests,* $\Delta$ $AIC_c = 0.0$ | | | |
| Intercept (Control) | 2.08 | 1.13 | 3.03 |
| Treatment | **1.48** | **0.16** | **2.79** |
| *Hatching success of nestlings; GLMM; n = 365 individuals from 69 nests,* $\Delta$ $AIC_c = 0.2$ | | | |
| Intercept (Control) | 2.16 | 1.19 | 3.13 |
| Treatment | **1.38** | **0.07** | **2.68** |
| Initial Brightness | **0.77** | **-0.04** | **1.58** |
| Treatment * Brightness | **-0.62** | **-2.01** | **0.76** |
| *Survival to day 12 for nestlings; binomial GLMM; n = 365 individuals from 69 nests,* $\Delta$ $AIC_c = 0.0$ | | | |
| Intercept (Control) | -0.05 | -0.95 | 0.86 |
| Treatment | **1.78** | **0.41** | **3.14** |
| Initial Brightness | **0.45** | **-0.37** | **1.28** |
| Treatment * Brightness | **0.70** | **-0.68** | **2.09** |
| *Survival to day 12 for nestlings; binomial GLMM; n = 365 individuals from 69 nests,* $\Delta$ $AIC_c = 1.3$ | | | |
| Intercept (Control) | -0.06 | -0.99 | 0.88 |
| Treatment | **1.74** | **0.34** | **3.14** |
| *Survival to fledge for nestlings; binomial GLMM; n = 365 individuals from 69 nests,* $\Delta$ $AIC_c = 0.0$ | | | |
| Intercept (Control) | **-0.89** | **-1.90** | **0.12** |
| Treatment | **1.34** | **-0.10** | **2.78** |
| Initial Brightness | **0.50** | **-0.37** | **1.37** |
| Treatment * Brightness | **0.57** | **-0.88** | **2.01** |
| *Survival to fledge for nestlings; binomial GLMM; n = 365 individuals from 69 nests,* $\Delta$ $AIC_c = 0.6$ | | | |
| Intercept (Control) | **-0.91** | **-1.94** | **0.13** |
| Treatment | **1.31** | **-0.15** | **2.78** |

**Table S16.** Details of simple models for associations between female feeding rate, glucose metabolism, and microbiome diversity versus nestling morphology and survival.

| Predictor | Response | Coefficient | Estimate | 2.5 % CI | 97.5 % CI |
|---|---|---|---|---|---|
| *Feeding Rate* | | | | | |
| | Mass | Intercept | 19.14 | 18.21 | 20.07 |
| | | Feeding Rate | 0.83 | -1.42 | 3.08 |
| | Wing length | Intercept | -0.02 | -0.27 | 0.23 |
| | | Feeding Rate | 0.01 | -0.59 | 0.62 |
| | Bill + head | Intercept | 0.03 | -0.20 | 0.26 |
| | | Feeding Rate | -0.03 | -0.59 | 0.53 |
| | # on Day 12 | Intercept | 3.58 | 3.12 | 4.03 |
| | | Feeding Rate | 1.49 | 0.35 | 2.63 |
| | # Fledged | Intercept | 2.86 | 2.40 | 3.33 |
| | | Feeding Rate | 2.10 | 0.94 | 3.26 |
| *Stress-Induced Glucose* | | | | | |
| | Mass | Intercept | 19.03 | 2.56 | 3.12 |
| | | Glucose | -0.79 | -1.68 | 0.11 |
| | Wing length | Intercept | -0.04 | -0.30 | 0.21 |
| | | Glucose | -0.08 | -0.33 | 0.17 |
| | Bill + head | Intercept | -0.01 | -0.25 | 0.22 |
| | | Glucose | -0.07 | -0.30 | 0.17 |
| | # on Day 12 | Intercept | 1.47 | -1.98 | 4.93 |
| | | Glucose | 0.01 | -0.01 | 0.02 |
| | # Fledged | Intercept | 2.31 | -1.33 | 5.95 |
| | | Glucose | 0.00 | -0.01 | 0.02 |
| *Microbiome Diversity* | | | | | |
| | Mass | Intercept | 14.78 | 11.54 | 18.03 |
| | | Simpson Diversity | 6.41 | 1.97 | 10.83 |
| | Wing length | Intercept | -0.15 | -1.06 | 0.76 |
| | | Simpson Diversity | 0.28 | -0.96 | 1.53 |
| | Bill + head | Intercept | 0.03 | -0.82 | 0.90 |
| | | Simpson Diversity | 0.04 | -1.14 | 1.21 |
| | # on Day 12 | Intercept | 3.20 | 1.40 | 5.00 |
| | | Simpson Diversity | 0.71 | -1.75 | 3.17 |
| | # Fledged | Intercept | 1.12 | -0.72 | 2.97 |
| | | Simpson Diversity | 2.62 | 0.09 | 5.15 |
